# Replay bursts coincide with activation of the default mode and parietal alpha network

**DOI:** 10.1101/2020.06.23.166645

**Authors:** Cameron Higgins, Yunzhe Liu, Diego Vidaurre, Zeb Kurth-Nelson, Ray Dolan, Timothy Behrens, Mark Woolrich

## Abstract

Our brains at rest spontaneously replay recently acquired information, but how this process is orchestrated to avoid interference with ongoing cognition is an open question. We investigated whether replay coincided with spontaneous patterns of whole brain activity. We found, in two separate datasets, that replay sequences were packaged into transient bursts occurring selectively during activation of the default mode network (DMN) and parietal alpha network. These networks were characterized by widespread synchronized oscillations coupled to increases in ripple band power, mechanisms that coordinate information flow between disparate cortical areas. Our data show a tight correspondence between two widely studied phenomena of neural physiology and suggest the DMN may coordinate replay bursts in a manner that minimizes interference with ongoing cognition.

## Main Text

A key mechanism by which the brain forms and stores new knowledge is through neural replay, whereby the patterns of neural activity associated with specific items are spontaneously reinstated in structured sweeps (*1*). These sweeps project to widespread regions of cortex (*2, 3*), with physiological signatures known as sharp wave ripples that have been described as the most synchronous events in the mammalian brain (*4*). Such spatially dispersed patterns of activity are often initiated during specific states, such as slow wave sleep, so as to preclude interference with ongoing wakeful processes. Replay was originally discovered during sleep, but is now known to occur also during wakefulness, particularly within periods of immobility and rest (*5–7*). An unanswered question in neuroscience is how the brain orchestrates these structured events in a manner that minimizes interference with ongoing cognition.

Neuroimaging studies have long highlighted that the brain in wakeful rest displays an intrinsic activity structure, cycling through a series of canonical resting state networks (RSNs) (*8–10*). One network that has drawn particular attention is the default mode network (DMN), a disparate set of brain regions that coactivate during ‘offline’ periods - such as between trials in the absence of specific tasks, and also during sleep (*11, 12*). The DMN has since been identified as correlating with a number of introspective cognitive states such as episodic memory and future oriented thought, suggesting a functional role during wakeful rest in mediating internally generated cognition and inhibiting bottom-up sensory processing (*13*). MEG resting state studies have established an ability to detect DMN activation (along with that of many other networks) with millisecond temporal resolution, demonstrating these networks activate transiently within specific spectrally defined modes (*10, 13–15*). An ability to measure replay noninvasively in humans has recently been demonstrated (*16, 17*), and this allowed us to investigate a potential link between replay and resting brain network activity.

Replay during slow wave sleep is associated with specific electrophysiological patterns; low frequency oscillations synchronize widespread regions of cortex, while high frequency sharp wave ripples propagate between hippocampus and cortical areas (*4, 18*). These widespread patterns appear integral to the effective function of replay in consolidating memories (*19*). In contrast, the wakeful brain displays a markedly different profile, with transient periods of synchronous activity interspersed with widespread desynchronisation associated with the distributed processing of parallel cognitive tasks (*10, 15*). Here we ask whether replay events, as detected by Liu et al. (*16*), are linked to specific changes in whole-brain neural activity, changes that might explain how wakeful replay could reinstate distributed cortical patterns from memory without interference from competing cognitive demands.

## Spontaneous replay coincides with activation of the Default Mode and Parietal Alpha networks

We investigated whether the replay events discovered by Liu et al. (*16*), each representing the rapid serial reactivation of learned stimulus representations (Fig. 1), coincided with specific macro patterns of resting brain network activity that have been studied widely in the literature (*10, 13–15*). The focus of our analysis was the same MEG scan data of Liu et al. (*16*), collected during resting periods of their experiment.

**Fig. 1.**
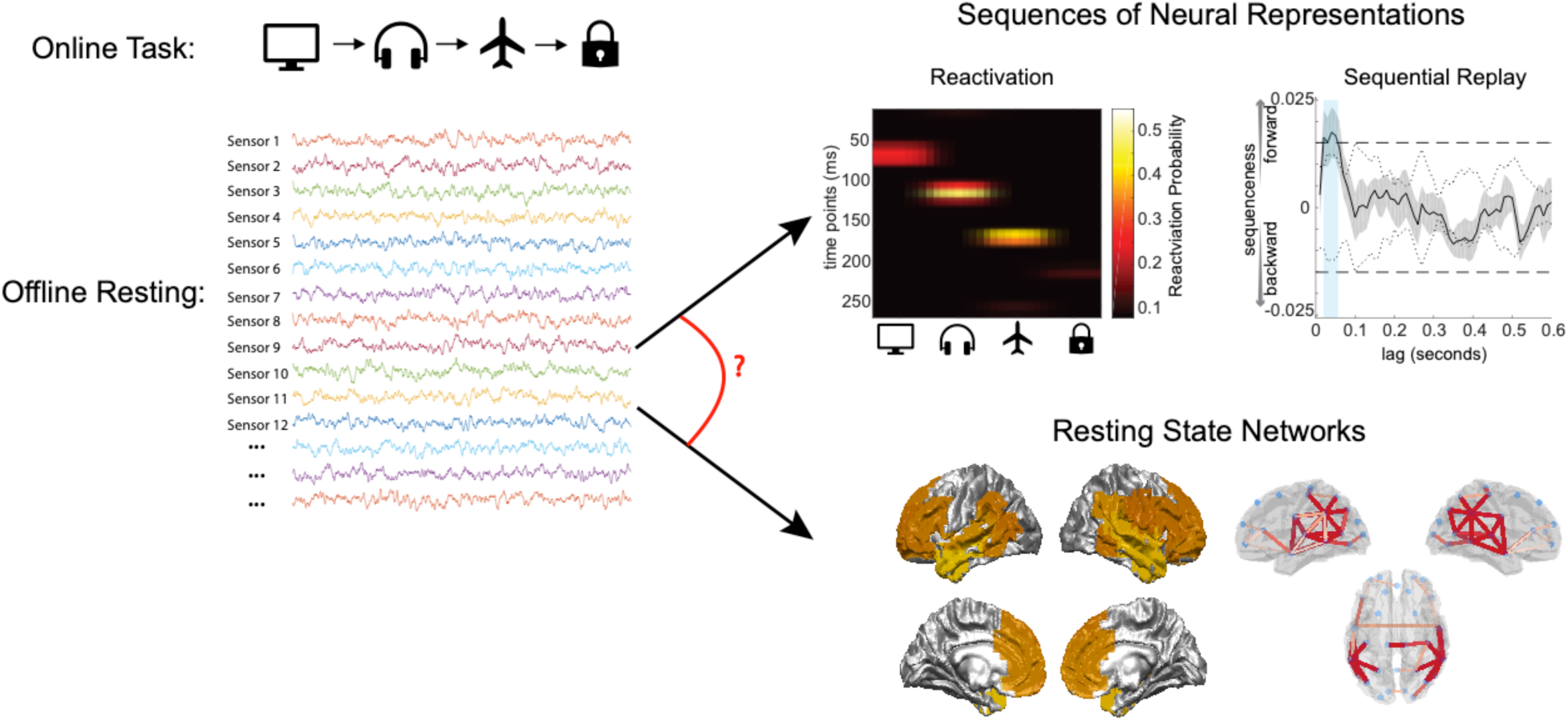
Task outline. The online task required participants to learn and remember a sequence of items. During the offline resting period, subjects were recorded passively with no immediate task; they were later tested for correct recall of the item sequence. The data from the offline resting period was analyzed in two ways: the first analysis detected replay using the methods of Liu et al. (*16*), using classifiers trained on the task items and identifying periods when they are reactivated in the specific sequence required by the task. The second analysis identified when specific resting state networks were activated by using an established model (*15*) to detect spontaneous patterns of spatial and spectral activity in the data. The objective was to determine whether the two measures were linked.

We first repeated the analysis of Liu et al. (*16*) to identify specific moments when replay occurred within this resting state data. Briefly, we trained multivariate classifiers to recognize each experimental stimulus, applied these classifiers to the resting data, and found the times when classifiers detected stimulus representations in rapid sequences played out in an order defined by the task (see Methods and Liu et al. (*16*)).

Next, in the same data, we determined which of a set of canonical resting state networks (RSN-states) were active at each point in time (see Methods and Fig. 1). We inferred 12 RSN-states in a data-driven way using an established Hidden Markov Model Framework (see methods and Vidaurre et al. (*15*)). These RSN-states were labelled according to a multidimensional scaling of their distances from each other (see Methods and Fig. S1); thus, RSN-states 1 and 12 represented opposing extremums of a single major axis of differentiation between networks. We then conducted an evoked response analysis to ask whether activation of the RSN-states was modulated around replay events.

As shown in Fig. 2A, a strong relationship emerged between replay onset and two RSN-states in particular, RSN-states 1 and 2. This relationship peaked at around t=0 - the exact time of replay-onset - but exhibited a decay at either side of this time, with a statistically significant association up to 0.5 seconds before and after each estimated replay-onset time (non-parametric cluster significance test, p<2e-4 for both RSN-states). In addition, a weaker relationship was also evident between RSN-states 3 and 4 and the observed replay times (p<2e-4 for both RSN-states).

**Fig. 2.**
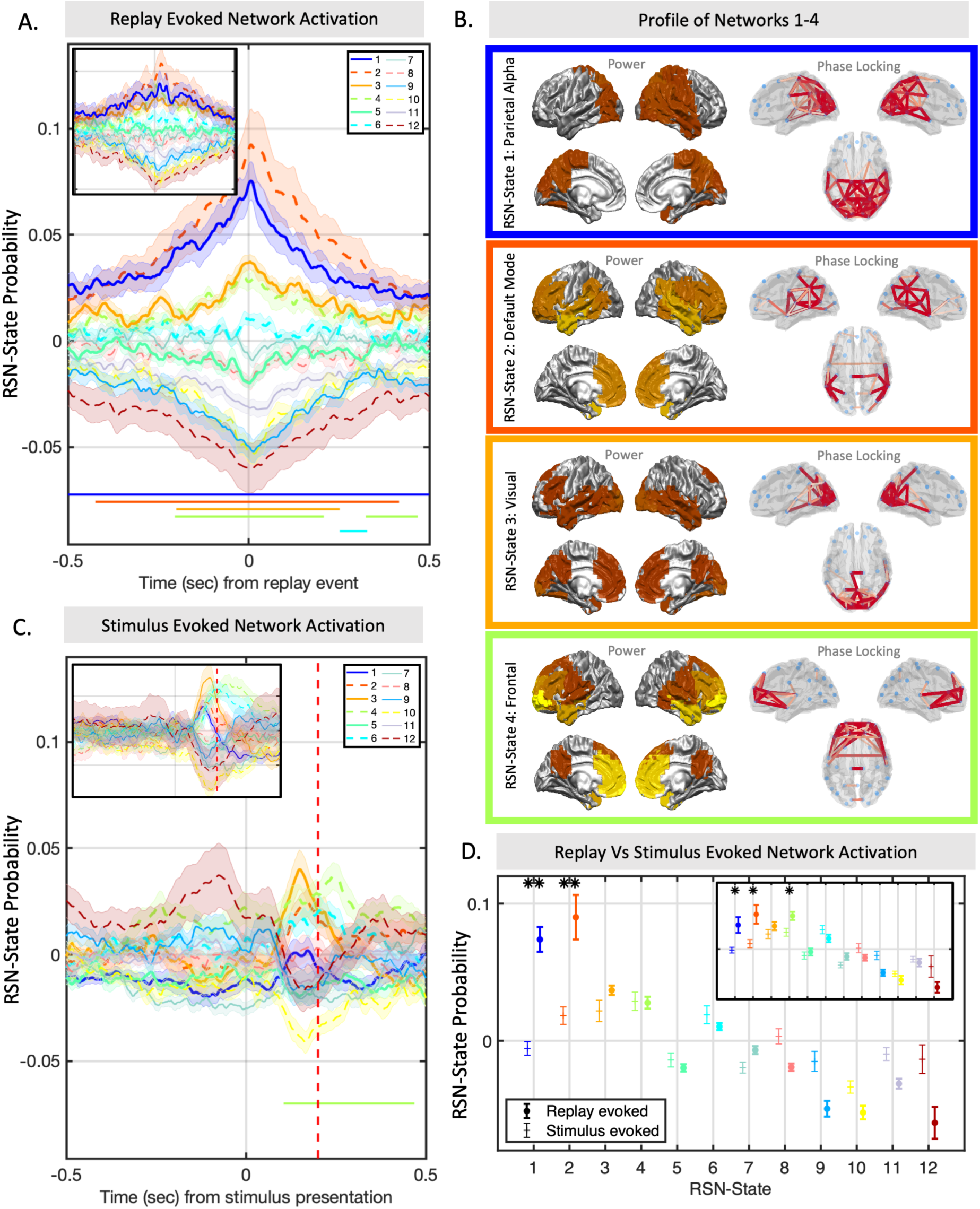
Replay coincides with activation of specific resting state networks. (A) A strong correlation exists between the replay times identified by Liu et al. (*16*) and resting network activity, namely RSN-States 1 and 2. This association peaks at t=0, representing the exact time of replay onset, but remains significant up to 0.5 seconds to either side of each event, showing that activity in either of these RSN-states is broadly predictive of replay. Significance bars show clusters where p<2e-4. Inset: Result of replication study on second dataset. (B) The broadband power and coherence networks that characterize each state identify RSN-states 1 and 2 as parietal alpha and default mode network respectively; RSN-states 3 and 4 correspond to activity in visual and frontal areas respectively. Power maps thresholded at 50%, phase locking networks thresholded with gaussian mixture model (see methods). (C) Fitting the same states to the original stimulus data on which the replay classifiers were trained identifies an overall network profile markedly different to that of the spontaneous replay events. Significance bars show clusters where p<2e-4. Inset: Result of replication study on second dataset. (D) Comparing directly the mean +/- standard error of the evoked state distribution at replay time and at the classifier training time identifies RSN-states 1 and 2 as significantly increased during spontaneous replay (multiple paired t tests). Single asterisk denotes p<0.05, double asterisk denotes p<5e-4. Inset: Result of replication study on second dataset.

Each RSN-state can be described by its distinct spectral power and phase locking profile; these profiles are summarized by averaging over frequency bands in Fig. 2B for RSN-states 1-4. This highlights that RSN-state 1 was associated with activity over parietal cortex; the equivalent network in simultaneous EEG-fMRI studies has been shown to be anticorrelated with the Dorsal Attention Network (DAN) (*20, 21*); thus the activation of RSN-state 1 corresponds to the DAN switching off. RSN-state 2 combines high power signals in frontal and temporal regions with coherent oscillations in lateral parietal cortex, regions that comprise the DMN. RSN-state 3 can be interpreted as activation of the visual cortex, and RSN-state 4 can be interpreted as activation of frontal cortical regions. The profiles of the remaining RSN-states are shown in Fig. S3; see Supplementary Text for more detail on how the MEG RSN-states relate to canonical resting state networks from fMRI.

The alignment of replay with specific RSN-states would not be surprising if some of these networks simply reflected the patterns of activity present during the original stimulus-encoding in the functional localizer data. Fig. 2C shows this is not the case; fitting the canonical RSNs to the original functional localizer data on which the classifiers were trained identified a markedly different relationship with the RSN-states compared with replay. No significant increase was observed to RSN-state 1 (p=0.88 at t=0.2sec), and RSN-state 2 showed a mild increase that was not significant after Bonferroni correction (p=0.006 at t=0.2 sec). Comparing directly the distribution of RSN-states evoked by replay and by the stimuli identified a clear differential distribution, with highly significant increases in the RSN-states 1 and 2 (Fig. 2D, one sided t-test p=8.9e-7 for RSN-state 1 and p=1.9e-4 for RSN-state 2). We conclude that the brain-wide patterns of resting RSN-state activity that are associated with replay, differ to those present during original stimulus-encoding. Instead, they are better characterized by activation of the default mode and parietal alpha RSN-states.

To ensure replicability of these results, the same analysis was conducted on a second independent study that examined replay data in MEG using a very similar but slightly amended paradigm (see (*16*)). We replicated the exact findings again, showing that RSN-states 1 and 2 had a strong association with replay (see Fig. 2A inset panels and Fig. S2; p<2e-4 for RSN-states 1 to 4). Neither RSN-states 1 or 2 displayed a significant increase in relation to the functional localizer data (p=0.67 for RSN-state 1 and p=0.14 for RSN-state 2). A paired t-test also confirmed these RSN-states were more strongly associated with replay than with the original training data (p=5.4e-3 for RSN-state 1 and p=9.1e-3 for RSN-state 2). Note that this is statistically weaker than what we observed in the first study, and neither result in the second study is significant after Bonferroni correction for multiple comparisons; however, considering the conservative nature of Bonferroni correction, we interpret it as a successful replication of the above reported findings.

### Transient replay bursts coincide with clusters of DMN and parietal alpha network activity

The correlation between replay and the RSN-states shown in Fig. 2B was maintained for over 0.5 seconds either side of each replay event. This is difficult to immediately reconcile with the highly transient nature of MEG RSN-states, which typically activate for less than 100msec. RSN-states 1 and 2 however have unique temporal profiles; replicating previous findings of distinct DMN dynamics (*14, 15*), when activated they endured for longer than any other RSN-states (Fig. 3B), and also quiesced for longer periods than any other RSN-states (Fig. 3C). Qualitative assessment suggested these state visits may cluster together over longer timescales, with clusters of DMN and parietal alpha network activity coinciding with bursts of replay events (Fig. 3A).

**Fig. 3.**
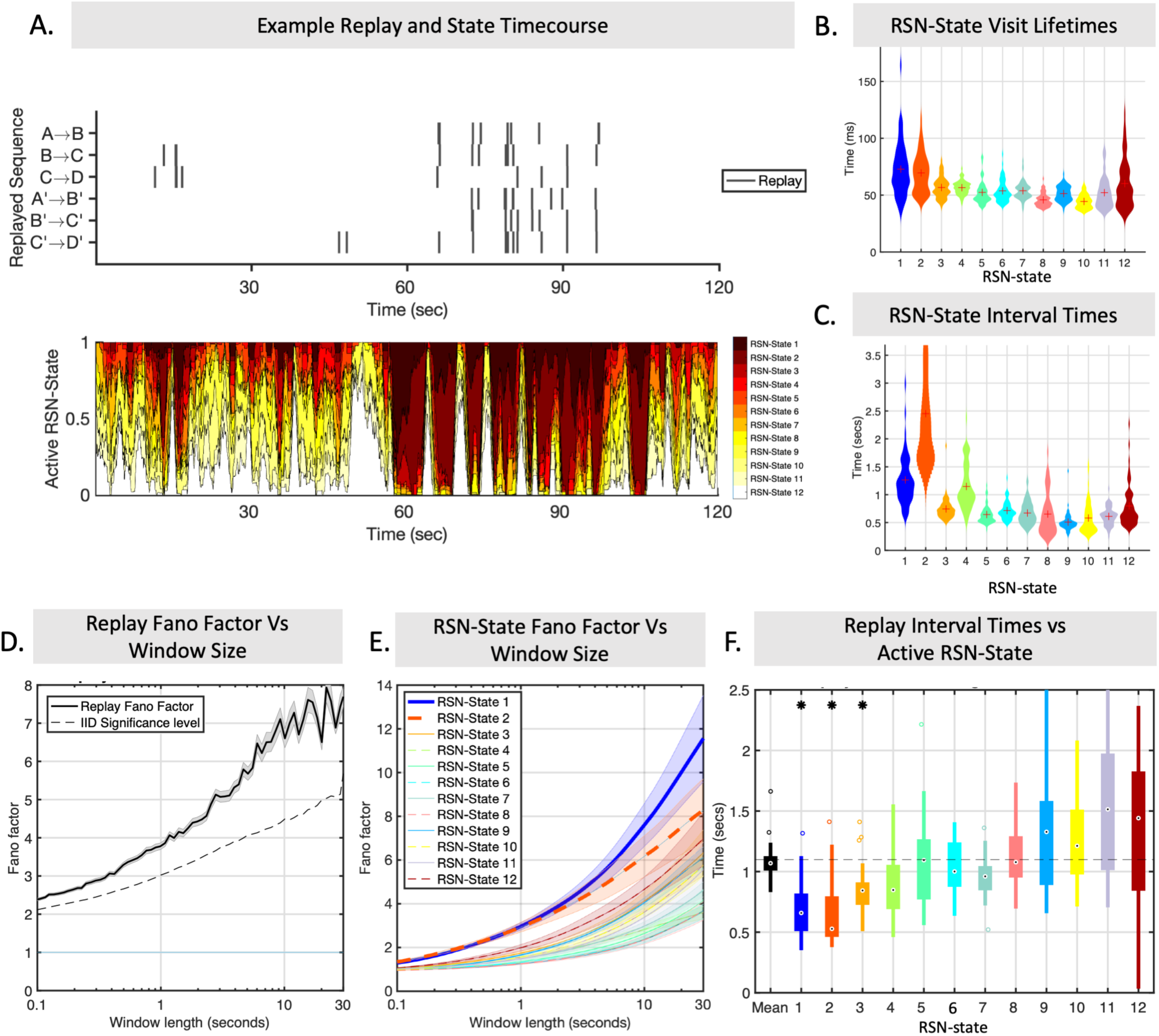
Replay and Resting State Networks share a common long-term temporal structure. (A) Example data showing the bursty nature of replay coinciding with relatively infrequent visits to the replay-associated RSN-states. For ease of visualization, lower panel plots the one second moving average RSN-state probability. (B) RSN-state visit lifetimes; the average time each RSN-state remains active when it is visited. RSN-states 1 and 2 display the longest lifetimes. (C) RSN-state interval times; the average time for each subject between visits to a particular RSN-state. RSN-states 1 and 2 have the longest interval times between visits. (D) Temporal irregularity can be quantified by looking at the Fano factor as a function of window size. Replay events show that this irregularity measure increases over longer timescales, displaying maximum temporal irregularity over windows of ten seconds or more. (E) This structure is replicated by the RSN-state activations, with RSN-states 1 and 2 displaying the most irregular patterns at long timescales. (F) This temporal structure is not just common but in fact coincides; replay events that occur during RSN-state 1 or 2 have significantly shorter periods, reflecting rapid bursty behavior during the infrequent state visits and long periods of quiescence outside of these.

To test whether replay events were concentrated into transient bursts, as suggested by (*16*), we first computed the Fano Factor over the time course of replay events; Fano factors equal to one correspond to a regular, non-bursting process, whereas Fano Factors greater than one corresponding to increasingly irregular bursting. As shown in Fig. 3D, the observed replay Fano Factor increased as a function of window size and exceeded one for all window sizes tested, showing that the occurrence of replay events was increasingly irregular over longer time periods. The bursting nature of replay events was further supported by the rejection of a broader null hypothesis that intervals between replay events were independent and identically distributed (p<1e-3; see Methods).

We conducted the same analysis on the time-courses of RSN-state occurrences (Fig. 3E). As with replay events, this showed that visits to different RSN-states were also increasingly irregular over longer time periods, but displayed a degree of irregularity that was not uniform over the different RSN-states (one-way ANOVA, p<2e-8 for all window sizes tested; this was significant after multiple comparison correction for RSN-states and number of windows). Notably, this was particularly pronounced for RSN-states 1 and 2 (two sample t-test p<6.4e-9 for RSN-state 1 and p<0.01 for RSN-state 2), the DMN and parietal alpha RSN-states that were most strongly correlated with replay. Thus, the RSN-states which were most inclined to cluster together into periods of increased intensity over long timescales were similarly the most correlated with replay.

We have found evidence that both the replay events and RSN-state visits display bursty behavior, resulting in clusters of intense activity interspersed with long periods of quiescence. In addition, we have evidence that replay events and RSN-state occurrence temporally coincide (cf. Fig. 2A). However, this alone does not necessarily mean that the bursting itself temporally coincides. To test whether this was the case, we computed the inter-replay interval time conditioned upon the active RSN-state at that time. We found that the interval to the next replay event was significantly determined by the currently active RSN-state (one way ANOVA, p=2e-9) such that when RSN-states 1 and 2 were active there were shorter intervals between replay events (p=2.6e-3 and p=5e-4 respectively). This suggests that replay events are packaged into bursts that occur selectively during clusters of intense DMN and Parietal Alpha RSN-state activity.

To assess reproducibility, we again replicated all results reported here on a second dataset of 22 subjects (see (*16*) and Methods); as shown in Fig. S4, replay Fano Factors again exceeded one, increased with window size, and exceeded non-parametric permutation test thresholds (p<1e-3); visits to RSN-states displayed a similar bursty profile, the degree of which was RSN-state dependent (p<1.5e-8, one-way ANOVA) with the Parietal Alpha and DMN RSN-states displaying the highest amount (two sample t-test: p<7e-6 for RSN-state 1, p<1.1e-4 for RSN-state 2); inter replay intervals were again significantly determined by active RSN-state (p=0.01, one-way ANOVA); RSN-state 1 was significantly associated with shorter intervals (p=6e-3), RSN-state 2 was trending in the same direction but not statistically significant (p=0.07).

### Replay coincides with distinct patterns of brain-wide highly synchronous activity

Having established a strong temporal association between replay and specific RSN-state activation, we next sought to characterize the nature of the brain-wide activity in the replay-associated RSN-states. For each RSN-state we calculated the spatial patterns of oscillatory power over all brain regions and the degree of synchronization (coherence) between all brain regions (as per (*15*); see Methods). Fig. 4A displays these averaged over brain regions, showing that the replay-associated RSN-states (RSN-states 1 and 2, and to a lesser extent, RSN-states 3 and 4) were associated with widespread increases in both the power and coherence of oscillatory activity compared with the other RSN-states (one way ANOVA for group-wise variation, p<1e-50 for both power and coherence; two sample t-tests, p<1e-50 for RSN-states 1-4 for both power and coherence).

**Fig. 4.**
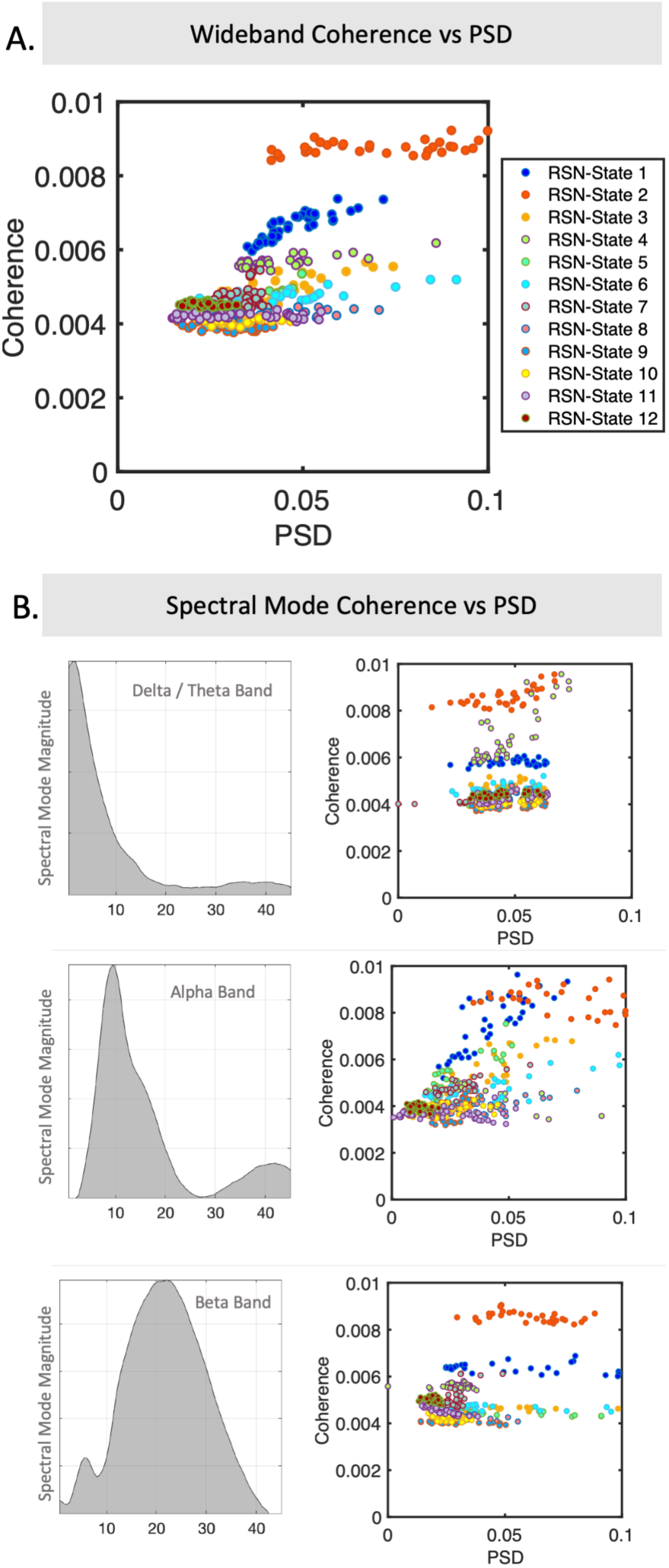
Replay associated RSN-states display specific spectral patterns of power and coherence. (A) Summarizing the frequency information in a single wideband plot, a scatter plot of the power spectral density (PSD) and coherence for each ROI (PSD is computed directly per ROI; coherence is taken as the sum of coherence values between that ROI and all others) as a function of active RSN-state identifies RSN-states 1 and 2 as having elevated coherence. (B) The data support a frequency decomposition into three modes that correspond to canonical delta/theta, alpha and beta bands (see methods). Scatter plots of each RSN-state’s PSD and coherence highlight that RSN-state 2 displays higher coherent activity across multiple frequency bands compared to all other RSN-states, whereas RSN-state 1 is more concentrated in the alpha band.

To better characterize the spatial distribution of activity in distinct frequency bands, we decomposed the spectral activity patterns using non-negative matrix factorization (*15*). This identified three prominent frequency modes reflecting activity in canonical frequency bands of delta/theta, alpha and beta bands (Fig. 4B). RSN-state 2, the DMN, showed a prominent elevation in network coherence across all three frequency bands compared to all other RSN-states; whereas RSN-state 1, the Parietal Alpha network, showed increased coherence in the alpha band compared to the other frequency bands. The two other replay-associated RSN-states were associated with activity concentrated in the alpha band (RSN-state 3) and delta/theta band (RSN-state 4).

We also used the RSN-state description of the resting state data to calculate the brain-wide patterns of oscillatory power and synchronization occurring specifically around replay events. Fig. 5 shows this first as time-frequency plots of power (Fig. 5A) and coherence (Fig. 5B) averaged over all brain regions, again highlighting a strong increase in power and coherence associated with each replay event. Fig. 5C shows the brain-wide patterns of oscillatory power (left) and synchronization (right) that occurred at the time of onset of the replay events. This highlights activity across both the frontal and parietal regions of the DMN, with activity in each brain region dissociated into two distinct frequency bands. Frontal nodes of the DMN, including the medial prefrontal cortex and temporal poles, were associated with coherent oscillations in a low delta / theta frequency band; parietal nodes, taken to include posterior cingulate and lateral parietal cortex, were associated with coherent oscillations in the alpha band.

**Fig. 5.**
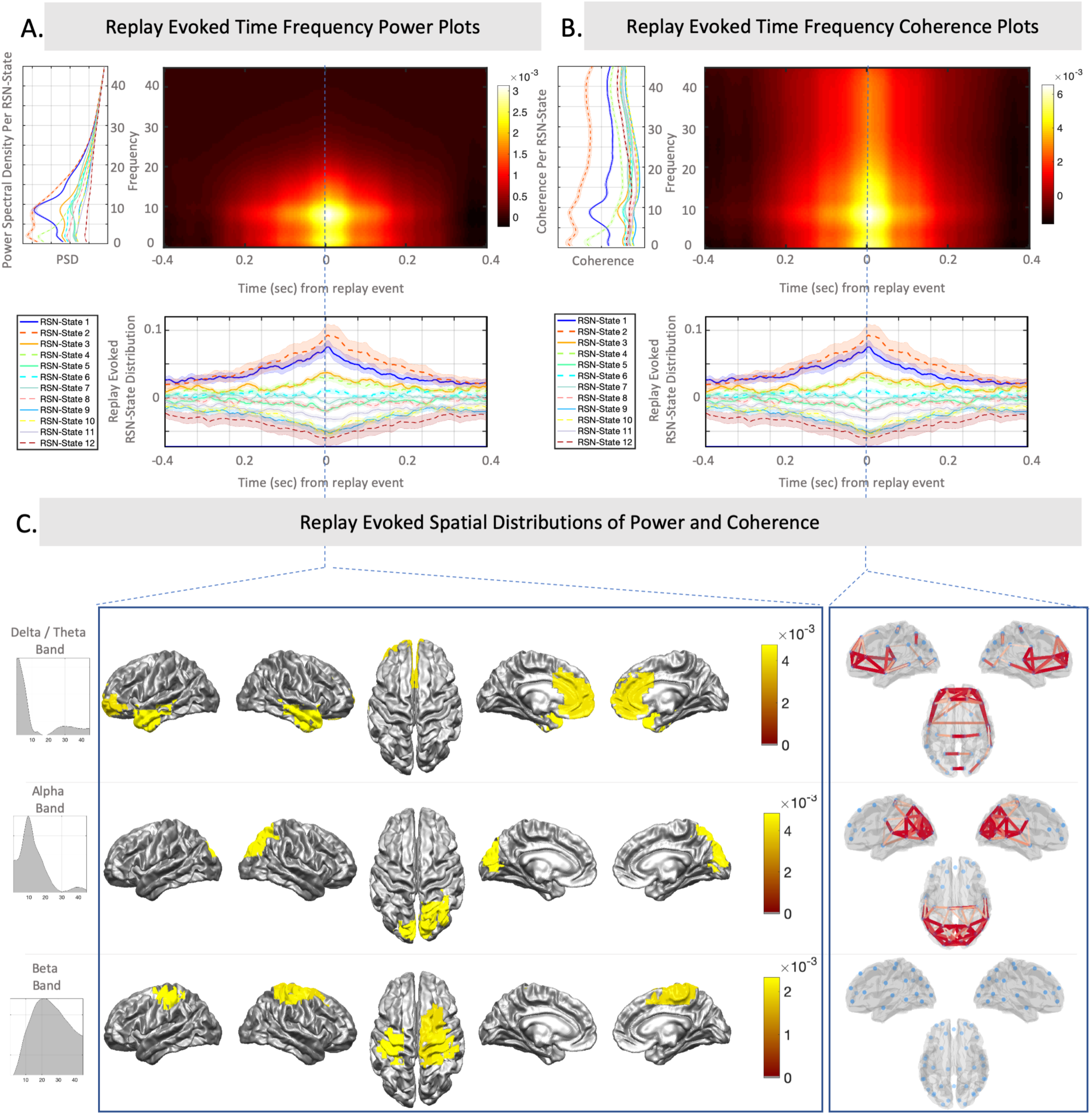
Replay associated brain activity is characterized by independent spatially and spectrally defined modes. (A) We can use the PSD estimates of each RSN-state (left) and the replay evoked RSN-state probabilities (lower) to reconstruct a time-frequency estimate of power spectral density around replay events, revealing a prominent peak in the alpha and delta /theta bands. (B) In addition to increases in power, the replay associated RSN-states show an increase in coherence across all frequencies, but especially in the alpha and delta/theta bands. (c) Plotting the spatial distribution of activity in the defined frequency modes at the time of replay identifies independent modes of coherent activity; a low frequency mode comprising frontal DMN and temporal areas, and an alpha frequency mode comprising parietal DMN regions and visual cortex. Additionally, some weaker levels of activity in the beta band are observed over motor areas, but network coherence patterns in this frequency band are not significant.

Again, in the interests of replicability, all these results were replicated on a second dataset of 22 subjects (see (*16*) and Methods), where all activation maps and spectro-spatial profiles were remarkably consistent; see Fig. S5.

### Sharp wave ripple-band power linked exclusively to activation of the DMN

Liu et al. found an association between the onset of replay and increases in high frequency (>100Hz) power, consistent with a model of sharp wave ripple activity that coincides with the detected replay events. Given the strong association between replay events and activity in DMN and Parietal Alpha networks in our work, we asked whether sharp wave ripple band activity might correlate more generally with activity in these networks. Crucially, our approach for estimating the RSN-states applied a low pass filter to the data with 45Hz cut-off frequency; so any correlation with power spectra above this cut-off can be interpreted as entirely independent of the original RSN-state estimation.

Fig. 6A plots the average high frequency power spectra over all periods when a given RSN-state is active; this demonstrates a significantly elevated power spectral density in the ripple-band associated exclusively with the DMN RSN-state (one way ANOVA p<2.5e-6 for each frequency band between 52 and 148Hz; p>0.05 if ANOVA excludes DMN). This relationship mimics that observed in the power spectral density averaged over 50msec windows around replay events (Fig. 6B), whilst accentuating the power increase in much higher frequencies. This suggests that the ripple band power bursts recorded by Liu et al., and taken to be synonymous with replay, may be a hallmark feature of DMN activation.

**Fig. 6.**
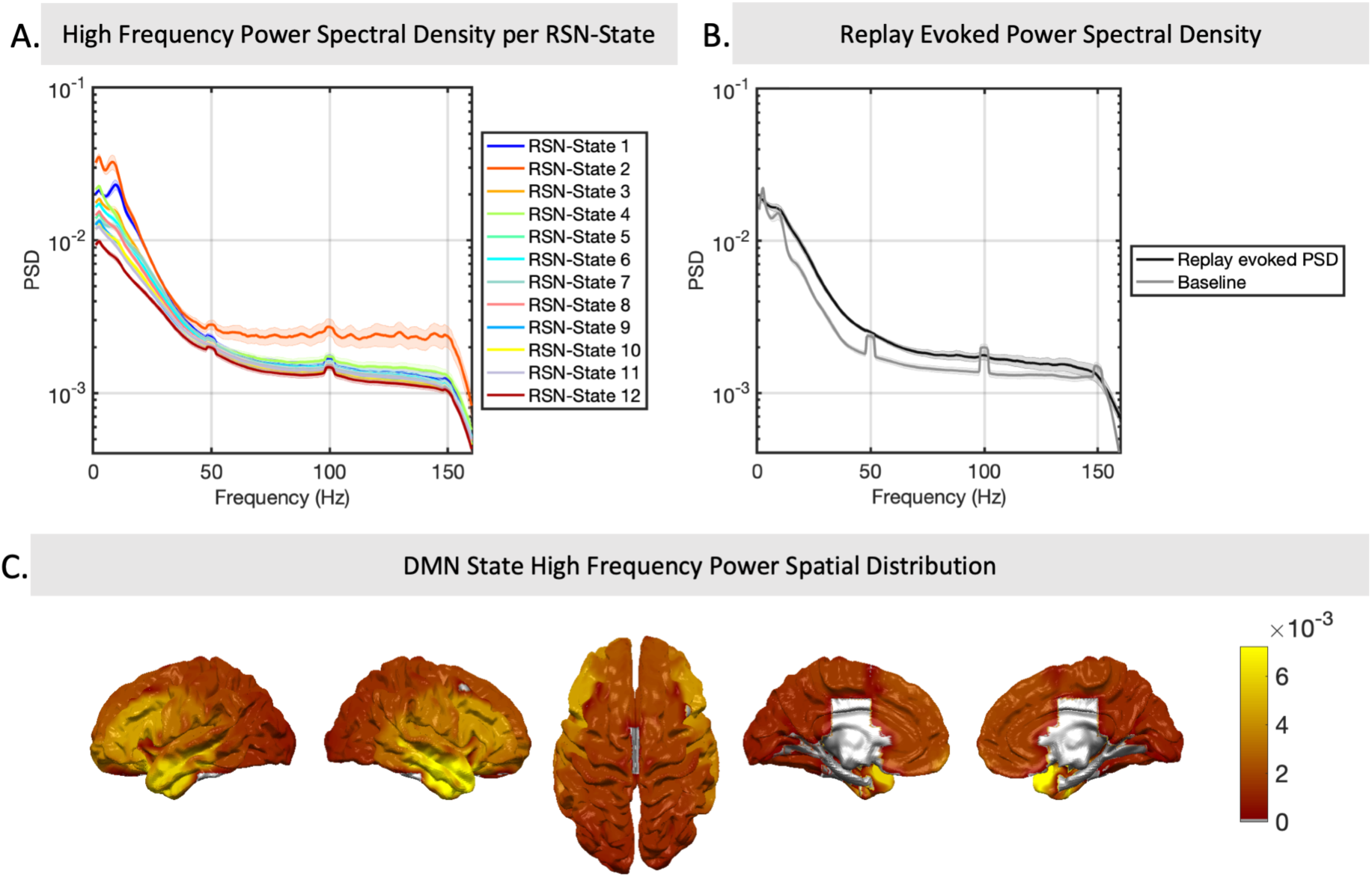
Both Replay and the DMN coincide with high frequency power increases in temporal areas. (A) Although the RSN-state model was originally fit to data filtered at 1-45Hz, we can still analyze whether the state timings correlate with specific patterns in frequency bands outside this range in the original data. This reveals a very strong association between RSN-state 2 and power in high frequencies – despite these high frequencies not having been originally included in the model. (B) Similarly, the onset of replay is associated with an increase in high frequency power relative to the global average. (C) Activity in this RSN-state and in this frequency band may originate in temporal cortices.

Plotting the spatial distribution of ripple-band power in the DMN RSN-state (Fig. 6C) shows a concentration of power in temporal cortex. While acknowledging the limitations of MEG in imaging deep sources, these results are nonetheless consistent with a model of hippocampal sharp wave ripple activity occurring selectively during activation of the DMN RSN-state.

Fig. S6 demonstrates the replication of these results on a second dataset of 22 subjects (see (*16*) and Methods) where the DMN was found to be significantly associated with elevated ripple band power (one way ANOVA; p<2.4e-6 for all frequencies between 52Hz and 148Hz; p>0.06 when DMN state omitted), with a spatial distribution of power concentrated over temporal cortices.

## Discussion

These results bridge two quite separate fields of enquiry in neuroscience. The study of replay has been predominantly characterized at the level of cellular connections and LFP oscillations in animals, while the study of resting brain networks has been largely the preserve of human neuroimaging. As such, the link we now establish between these has the potential to extend not only our understanding of replay but also our understanding of the functional role played by human resting brain networks. We have looked for *sequential* reactivations consistent with the task sequence, but similar analyses work if we simply consider *all* reactivations (see Supplementary Text and Fig. S9). It has been established in these datasets that reactivations are more likely to occur in relevant sequences than control sequences *(16)*. However, because reactivations are bursty, we cannot make selective claims about sequential reactivations here. Increases in the density of *sequential* reactivations coincide with increases in the density of *all* reactivations, and these times align to periods of DMN and parietal alpha activity.

Both the DMN and parietal alpha activity have parallel histories in the scientific literature, initially interpreted as reflecting idling or default patterns of activity and only subsequently understood to have functional roles supporting attention and cognition (*22, 23*). The DMN in particular has since been associated with a role broadly defined as reflecting internally oriented cognition, encompassing functions like episodic memory and future oriented thought (*23, 24*). But in the same way that the brain uses sleep to replay past experience and consolidate memories, our results suggest that healthy waking brain activity may undergo periods of heightened DMN and alpha activity to perform the same function alongside ongoing cognition. Given our more refined mechanistic understanding of the role of replay, our new findings could extend our interpretation of the functional relevance of the DMN. Replay itself is fundamentally understood as a mechanism for memory consolidation, but has also been proposed to have more expansive roles in building cognitive maps, preparing neural structures for learning (preplay) and transferring knowledge from hippocampus to cortex (*7, 16, 25*). This suggests a more expansive role for the DMN in executive control, with the regular transient activations of the DMN associated with building and maintaining stable representations of recently acquired information.

Replay occurs in the highest intensity during slow wave sleep, when large scale synchronized oscillations provide an environment conducive to the large scale propagation of sharp wave ripples (*26*). Notably, the RSN-states that correlate with replay in our study are characterized by large increases in oscillatory coherence, itself hypothesized to support integration of signals from disparate regions of cortex. The DMN itself appears to be preserved at least into light sleep stages (*27–29*), with further evidence that the DMN correlates directly with sharp wave ripples under light anesthesia (*30*). Our findings appear to reflect both these phenomena – low frequency coherent oscillations and high frequency sharp wave ripples – coinciding directly with estimated replay events during an awake state. Importantly the high frequency band power increases we see alongside DMN activation appear much more broadband than has been established by studies of human sharp wave ripple activity using invasive methods (*31, 32*). We do not have a good explanation for this at present, but it is notable that relative power increases on short timescales aligned to the replay events themselves, in the same datasets, produces a more narrow-band high-frequency spectrum (*16*).

The role of the parietal alpha network is perhaps more readily understood through the unique requirements of awake state replay. Our results are based on spontaneous replay of visual stimuli representations, raising the problem of how the brain could reinstate these items without interference from ongoing perception. Alpha oscillations are widely interpreted as an inhibitory signal that acts to gate irrelevant stimuli from active processing (*33–35*). One possibility is that strong alpha activity may combine with DMN activation to inhibit bottom-up sensory perception during inward oriented attention, thereby supporting replay of items from memory. This could provide a crucial mechanism for how replay plays out without interference from competing sensory inputs during the awake state.

The temporal profile of replay activation that we have characterized may also explain replay related signals across different recording modalities. In particular, fMRI studies have shown reliable behavioral correlations between the reinstatement of BOLD traces associated with experimental stimuli and subsequent task performance (*36–38*). This has supported an interpretation of the BOLD signal as a reflection of cellular replay despite the disparity of timescales between cellular replay events (known to be temporally compressed on the order of milliseconds), and the hemodynamic response (assumed to reflect sustained activations on the order of seconds). However, our results characterize replay as occurring in transient bursts of high intensity, interspersed by long periods of quiescence. Such a temporal profile may bridge this disparity of timescales and explain how a hemodynamic signal could arise from clusters of replay bursts in quick succession.

Furthermore, our results can help to bridge the understanding of RSNs studied in electrophysiology and fMRI. A longstanding challenge in unifying findings across modalities has been to understand how the BOLD response relates to activity in canonical frequency bands. Activity in the gamma band (>30Hz) has consistently shown a strong correlation with a subsequent BOLD signal (*39, 40*), however a more complex relationship emerges between the BOLD signal and activity in lower frequency bands, in which electrophysiological RSNs are largely defined (*10, 20*). It has now been shown that different RSN-states show markedly different hemodynamic profiles; in particular, the DMN and DAN evoke BOLD signals that are both opposed in polarity and distinct in their temporal decay profile (*21*). But if the DMN state visits cluster together in time whilst being linked to high frequency power increases, as our results indicate, then this may help to explain these distinct profiles and provide a key bridge between the understanding of RSNs recorded across distinct modalities.

Finally, our results suggest measures that could potentially serve as non-invasive proxy measures of replay, potentially opening the door to a broader set of replay experimental paradigms. Until very recently, replay has been predominantly studied in animal models using spatial navigation paradigms, due to the necessity of highly invasive electrophysiology in order to detect replay and the sophisticated understanding of the entorhinal-hippocampal spatial navigation systems. The development of methods to detect reactivation (*38*) and replay (*16, 17, 41*) in humans noninvasively has enabled experiments that test how these theories generalize to non-spatial domains and other abstract cognitive tasks that are unique to human neuroscience. However, these methods require demanding experimental designs and cognitive paradigms. Our results broaden the range of tools further, providing additional non-invasive measures that could provide an estimate of aggregate replay activity under simple experimental conditions (rest) that can be potentially be studied in large populations or patient groups.

Overall, our results highlight an important link between two influential concepts in modern neuroscience; the study of replay and the study of resting state networks, and provide a connection between noninvasive human imaging studies and invasive cellular physiology.

## Acknowledgments

we are grateful to Andrew Quinn for his guidance on data processing, beamforming and visualization code; and to Jiewon Kang for her preliminary investigations of high frequency oscillations.

## Funding

The Wellcome Centre for Integrative Neuroimaging is supported by core funding from the Wellcome Trust (203139/Z/16/Z). MWW’s research is supported by the NIHR Oxford Health Biomedical Research Centre, by the Wellcome Trust (106183/Z/14/Z, 215573/Z/19/Z), and by the New Therapeutics in Alzheimer’s Diseases (NTAD) study supported by UK MRC and the Dementia Platform UK.

## Author contributions

Y.L., C.H., R.D., Z.K.-N., M.W. and T.E.J.B. conceptualized the research goals; M.W. and T.E.J.B. supervised the investigation; D.V., M.W and C.H. developed the methodology and software implementation; Y.L. collected and curated the data and assisted with formal data analysis; C.H. formally analyzed the data and wrote the initial draft; all authors reviewed and edited the manuscript;

## Competing interests

Z.K.-N. is employed by DeepMind Technologies Limited;

## Data and materials availability

All data and software code used to implement the analysis presented is freely available upon request to the lead author.

## References

1. M. A. Wilson, B. L. McNaughton, Reactivation of hippocampal ensemble memories during sleep. Science. 265, 676–679 (1994).

2. Y. L. Qin, B. L. McNaughton, W. E. Skaggs, C. A. Barnes, Memory reprocessing in corticocortical and hippocampocortical neuronal ensembles. The hippocampal and parietal foundations of spatial cognition. 352, 305–319 (1997).

3. D. Ji, M. A. Wilson, Coordinated memory replay in the visual cortex and hippocampus during sleep. Nature Neuroscience. 10, 100–107 (2007).

4. G. Buzsáki, Hippocampal sharp wave-ripple: A cognitive biomarker for episodic memory and planning. Hippocampus. 25, 1073–188 (2015).

5. H. S. Kudrimoti, C. A. Barnes, B. L. McNaughton, Reactivation of hippocampal cell assemblies: effects of behavioral state, experience, and EEG dynamics. The Journal of Neuroscience. 19, 4090–4101 (1999).

6. M. F. Carr, S. P. Jadhav, L. M. Frank, Hippocampal replay in the awake state: A potential substrate for memory consolidation and retrieval. Nature Neuroscience. 14, 147–153 (2011).

7. D. J. Foster, Replay comes of age. Annual Review of Neuroscience. 40, 581–602 (2017).

8. B. Biswal, F. Z. Yetkin, V. M. Haughton, J. S. Hyde, Functional connectivity in the motor cortex of resting human brain using echo-planar mri. Magnetic Resonance in Medicine. 34, 537–541 (1995).

9. M. D. Fox, M. E. Raichle, Spontaneous fluctuations in brain activity observed with functional magnetic resonance imaging. Nature Reviews Neuroscience. 8, 700–11 (2007).

10. M. J. Brookes, M. Woolrich, H. Luckhoo, D. Price, J. R. Hale, M. C. Stephenson, G. R. Barnes, S. M. Smith, P. G. Morris, Investigating the electrophysiological basis of resting state networks using magnetoencephalography. Proceedings of the National Academy of Sciences. 108, 16783–16788 (2011).

11. M. E. Raichle, A. M. Macleod, A. Z. Snyder, W. J. Powers, D. A. Gusnard, G. L. Shulman, A default mode of brain function. Proceedings of the National Academy of Sciences. 98, 676–682 (2001).

12. M. Fukunaga, S. G. Horovitz, P. van Gelderen, J. A. de Zwart, J. M. Jansma, V. N. Ikonomidou, R. Chu, R. H. R. Deckers, D. A. Leopold, J. H. Duyn, Large-amplitude, spatially correlated fluctuations in BOLD fMRI signals during extended rest and early sleep stages. Magnetic Resonance Imaging. 24, 979–992 (2006).

13. F. de Pasquale, S. della Penna, A. Z. Snyder, C. Lewis, D. Mantini, L. Marzetti, P. Belardinelli, L. Ciancetta, V. Pizzella, G. L. Romani, M. Corbetta, Temporal dynamics of spontaneous MEG activity in brain networks. Proceedings of the National Academy of Sciences. 107, 6040–6045 (2010).

14. A. P. Baker, M. J. Brookes, I. A. Rezek, S. M. Smith, T.E.J. Behrens, P. J. P. Smith, M. Woolrich, Fast transient networks in spontaneous human brain activity. eLife. 2014, 1–18 (2014).

15. D. Vidaurre, L. T. Hunt, A. J. Quinn, B. A. E. Hunt, M. J. Brookes, A. C. Nobre, M. W. Woolrich, Spontaneous cortical activity transiently organises into frequency specific phase-coupling networks. Nature Communications. 9 (2018), doi:10.1038/s41467-018-05316-z.

16. Y. Liu, R. J. Dolan, Z. Kurth-Nelson, T. E. J. Behrens, Human replay spontaneously reorganises experience. Cell. 178, 1–13 (2019).

17. Z. Kurth-Nelson, M. Economides, R. J. Dolan, P. Dayan, Fast sequences of non-spatial state representations in humans. Neuron. 91, 194–204 (2016).

18. J. J. Chrobak, G. Buzsáki, High-frequency oscillations in the output networks of the hippocampal-entorhinal axis of the freely behaving rat. Journal of Neuroscience. 16, 3056–3066 (1996).

19. G. Buzsaki, Memory consolidation during sleep: a neurophysiological perspective. Journal of sleep research. 7 Suppl 1, 17–23 (1998).

20. D. Mantini, M. G. Perrucci, C. del Gratta, G. L. Romani, M. Corbetta, Electrophysiological signatures of resting state networks in the human brain. Proceedings of the National Academy of Sciences. 104, 13170–13175 (2007).

21. T. Sitnikova, J. W. Hughes, C. M. Howard, K. A. Stephens, M. Woolrich, D. H. Salat, bioRxiv, in press, doi:10.1101/2020.05.05.079749.

22. W. J. Ray, H. W. Cole, EEG alpha activity reflects attentional demands, and beta activity reflects emotional and cognitive processes. Science. 228, 750–752 (1985).

23. M. E. Raichle, The brain’s default mode network. Annual review of neuroscience. 38, 433–47 (2015).

24. J. R. Andrews-Hanna, J. S. Reidler, C. Huang, R. L. Buckner, Evidence for the default network’s role in spontaneous cognition. Journal of Neurophysiology. 104, 322–335 (2010).

25. G. Dragoi, S. Tonegawa, Preplay of future place cell sequences by hippocampal cellular assemblies. Nature. 469, 397–401 (2011).

26. A. Sirota, J. Csicsvari, D. Buhl, G. Buzsáki, Communication between neocortex and hippocampus during sleep in rodents. Proceedings of the National Academy of Sciences of the United States of America. 100, 2065–2069 (2003).

27. M. D. Greicius, V. Kiviniemi, O. Tervonen, V. Vainionpää, A. L. Reiss, V. Menon, Persistent default-mode network connectivity during light sedation. Human brain mapping. 29, 839–847 (2008).

28. L. J. Larson-Prior, J. M. Zempel, T. S. Nolan, F. W. Prior, A. Z. Snyder, M. E. Raichle, Cortical network functional connectivity in the descent to sleep. Proceedings of the National Academy of Sciences. 106, 4489–4494 (2009).

29. P. G. Sämann, R. Wehrle, D. Hoehn, V. I. Spoormaker, H. Peters, C. Tully, F. Holsboer, M. Czisch, Development of the brain’s default mode network from wakefulness to slow wave sleep. Cerebral Cortex. 21, 2082–2093 (2011).

30. R. Kaplan, M. H. Adhikari, R. Hindriks, D. Mantini, Y. Murayama, N. K. Logothetis, G. Deco, Hippocampal sharp-wave ripples influence selective activation of the default mode network. Current Biology. 26, 686–691 (2016).

31. N. Axmacher, C. E. Elger, J. Fell, Ripples in the medial temporal lobe are relevant for human memory consolidation. Brain. 131, 1806–1817 (2008).

32. A. P. Vaz, S. K. Inati, N. Brunel, K. A. Zaghloul, Coupled ripple oscillations between the medial temporal lobe and neocortex retrieve human memory. Science. 363, 975–978 (2019).

33. O. Jensen, A. Mazaheri, Shaping functional architecture by oscillatory alpha activity: gating by inhibition. Frontiers in human neuroscience. 4, 186 (2010).

34. T. A. Rihs, C. M. Michel, G. Thut, Mechanisms of selective inhibition in visual spatial attention are indexed by α-band EEG synchronization. European Journal of Neuroscience. 25, 603–610 (2007).

35. J. J. Foxe, A. C. Snyder, The role of alpha-band brain oscillations as a sensory suppression mechanism during selective attention. Frontiers in Psychology. 2, 1–13 (2011).

36. I. Momennejad, A. Ross Otto, N. D. Daw, K. A. Norman, Offline replay supports planning: fMRI evidence from reward revaluation. eLife, 7, e32548 (2018).

37. L. Deuker, J. Olligs, J. Fell, T. A. Kranz, F. Mormann, C. Montag, M. Reuter, C. E. Elger, N. Axmacher, Memory consolidation by replay of stimulus-specific neural activity. Journal of Neuroscience. 33, 19373–19383 (2013).

38. A. Tambini, L. Davachi, Awake reactivation of prior experiences consolidates memories and biases cognition. Trends in Cognitive Sciences. 23, 876–890 (2019).

39. J. Niessing, B. Ebisch, K. E. Schmidt, M. Niessing, W. Singer, R. A. W. Galuske, Hemodynamic signals correlate tightly with synchronized gamma oscillations. Science. 309, 948–951 (2005).

40. Y. Nir, L. Fisch, R. Mukamel, H. Gelbard-Sagiv, A. Arieli, I. Fried, R. Malach, Coupling between neuronal firing rate, gamma LFP, and BOLD fMRI is related to interneuronal correlations. Current Biology. 17, 1275–1285 (2007).

41. N. W. Schuck, Y. Niv, Sequential replay of nonspatial task states in the human hippocampus. Science. 364, 1–9 (2019).

42. B. A. E. Hunt, P. K. Tewarie, O. E. Mougin, N. Geades, D. K. Jones, K. D. Singh, P. G. Morris, P. A. Gowland, M. J. Brookes, Relationships between cortical myeloarchitecture and electrophysiological networks. Proceedings of the National Academy of Sciences. 113, 13510–13515 (2016).

43. M. X. Huang, J. C. Mosher, R. M. Leahy, A sensor-weighted overlapping-sphere head model and exhaustive head model comparison for MEG. Physics in Medicine and Biology. 44, 423–440 (1999).

44. G. L. Colclough, M. W. Woolrich, P. K. Tewarie, M. J. Brookes, A. J. Quinn, S. M. Smith, How reliable are MEG resting-state connectivity metrics? NeuroImage. 138, 284–293 (2016).

45. G. L. Colclough, M. J. Brookes, S. M. Smith, M. W. Woolrich, A symmetric multivariate leakage correction for MEG connectomes. NeuroImage. 117, 439–448 (2015).

46. D. Vidaurre, R. Abeysuriya, R. Becker, A. J. Quinn, F. Alfaro-Almagro, S. M. Smith, M. W. Woolrich, Discovering dynamic brain networks from big data in rest and task. NeuroImage. 180, 646–656 (2018).

47. D. Vidaurre, A. J. Quinn, A. P. Baker, D. Dupret, A. Tejero-Cantero, M. W. Woolrich, Spectrally resolved fast transient brain states in electrophysiological data. NeuroImage. 126, 81–95 (2016).

48. M. C. Teich, R. G. Turcott, R. M. Siegel, Temporal correlation in cat striate-cortex neutral spike trains. IEEE Engineering in Medicine and Biology. 15, 79–87 (1996).

49. M. Corbetta, G. L. Shulman, Control of goal-directed and stimulus-driven attention in the brain. Nature Reviews Neuroscience. 3, 201–215 (2002).

