## Supplementary Info for "Replay bursts coincide with activation of the default mode and parietal alpha network"

### Supplementary Information

**This PDF file includes:**

Materials and Methods
Supplementary Text
Figs. S1 to S9

#### Materials and Methods

Datasets

We make reference throughout this publication to three separate datasets; we will refer to these as Dataset A, the replay data collected by Liu et al that forms the main focus of analysis in this paper; Dataset B, a secondary replay data set that was used to prove replication of the analysis conducted on dataset A, which is referred to throughout the paper and is the focus of Fig. S2, S4, S5 and S6; and Dataset C, an established, larger dataset comprising resting state MEG data that we used to train the canonical resting state networks.

Dataset A, the primary replay dataset, was acquired from 25 participants (aged 19-34, mean age 24.89); four subjects were then excluded due to large motion artefacts or missing trigger information, leaving 21 subjects for the analysis conducted. Dataset B, the replication dataset, was acquired from 26 participants (aged 19-34, mean age 25.48); four participants were later excluded due to motion artefacts or failure to complete the task, leaving 22 subjects for the analysis conducted. In both Dataset A and B, MEG scans were acquired at 600 samples/second on a 275 channel CTF MEG system. All participants signed written consent in advance; ethical approval for the experiment was obtained from the Research Ethics Committee at University College London under ethics number 9929/002.

Dataset C, the RSN-state training dataset, comprised resting state MEG data from a larger group of subjects that has previously been used to characterize MEG resting state network dynamics *(15,42)*. This study acquired resting state MEG scans and structural MRI scans for 55 participants (mean age 26.5 years, maximum age 48 years, minimum age 18 years). MEG data was acquired at 1200Hz using a 275 channel CTF MEG system operating at third order synthetic gradiometry configuration; MRI data were acquired using a Phillips Achieva 7T system. MRI data, used only for the purpose of MEG coregistration, were acquired using a Philips Achieva 7T scanner. All participants gave written informed consent and ethical approval was granted by the University of Nottingham Medical School Research Ethics Committee.

Data Preprocessing

Across the three datasets and the two analysis pipelines of figure 1b we sought to minimize differences in preprocessing however minor deviations were necessary. The replay identification conducted by Liu et. al. (Fig. 1) filtered sensor space MEG data to a pass band of 0.5 to 50 Hz; data were downsampled to 100 samples / second. All analysis reported by Liu et. al. *(16)* in this paper was conducted in sensor space; for further details see *(16)*. The Resting State Network analysis pipeline filtered sensor space data to a pass band of 1 to 45 Hz and downsampled data to 250 samples / second. This slightly amended filter passband was to ensure RSN-state dynamics were not driven by low frequency sensor drift effects or mainline power noise effects, which the HMM modelling approach is more sensitive to compared to the replay identification methods introduced by Liu et. al. *(16)*. Similarly, the higher sampling rate was to ensure sufficient resolution of the time embedding to ensure good estimation of spectral content for each RSN-state definition. The only exception to this was for the final analysis of spectral content in higher frequencies (Fig. 6), for which we returned to the raw data to utilize the highest frequency information available; this data had been acquired at 600 samples / second and low pass filtered with cutoff frequency 160Hz.

Source Space Reconstruction

One of the motivations for using Dataset C to train the resting state networks was to ensure the highest possible confidence and replicability of the anatomical distributions of activity unique to each RSN-state. Given the limited spatial resolution of MEG, confidence can be increased through the use of larger datasets which have been coregistered using MRI acquired structural scans. MRI scans were not acquired for either dataset A or dataset B, motivating our focus on dataset C to determine high confidence spatial topographies of each RSN-state.

Dataset C was coregistered to MRI structural information using a multiple local sphere forward model *(43)*. Datasets A and B, which did not have associated MRI structural information, were coregistered using fiducial markers. All data then followed a common pipeline of analysis thereafter, with source space reconstruction performed using an LCM-V beamformer projecting to an 8mm MNI grid. This grid was then parcellated into 38 anatomically defined Regions of Interest (ROIs) derived from an independent component analysis of fMRI resting state data from the Human Connectome Project *(44)*. Source leakage was then corrected for by orthogonalization as outlined in *(45)*.

Replay Experimental protocol

For full details of the experimental protocol, readers are directed to Liu et. al. *(16)*; key details only are summarized here. In Dataset A, each participant attended two days, the first for learning the structure of the task and the second for completing the task while undergoing MEG scanning. The task was based around 8 visual stimuli, and participant's objective was to correctly unshuffle these into the two correct four item-long sequences (eg, A->B->C->D and A' -> B' -> C' -> D'), using a set of unshuffling rules learned on the first day of the experiment. Once in the scanner, participants observed multiple presentations of the novel visual stimuli in a randomized order to act as training data for the multivariate classifiers (the Functional Localizer data. They were then presented with the visual stimuli in the shuffled order from which they could infer the correct sequence. Participants then underwent the first 5 minute resting state scan; one of the two four-item sequences was paired with a reward; participants underwent a second resting state scan, and were tested on their correct recall of the item sequences. All analysis in this paper focusses only on the data from the two resting state scans within this overall experiment. Dataset B used a very similar but slightly amended paradigm; the shuffling rule that participants learned was different, as was the ordering of individual blocks (see *(16)*).

Replay Detection

This paper builds on the results of *(16)*, using their computed replay onset timings. For clarity we outline their methods to detect replay here, but direct readers to the original publication for further details. For each of the eight visual stimuli, sparse logistic regression classifiers were trained using L1 regularization on the functional localizer data to identify the neural patterns associated with each visual stimulus. To select an appropriate timepoint on which to train the classifiers, the authors used cross validation over the functional localizer data to plot the overall classification accuracy; this identified the timepoint 200msec following stimulus presentation to be the timepoint corresponding to the highest classification accuracy averaged over all stimuli, trials and subjects. The trained classifiers associated with this timepoint were then fit to the resting state data, producing eight time series that represented the probability of reinstatement of the activity patterns associated with each visual stimulus:

$$A_t = \sigma(X_t \beta_A)$$

$$B_t = \sigma(X_t \beta_B)$$
...

Where $X_t$ is the recorded resting state data at time $t$; $\beta_i$ is the sparse classifier coefficients associated with the $i$th visual stimulus; $A_t$ is the probability of reinstatement of the patterns associated with stimulus $A$ at time $t$ in the resting state scan; and $\sigma(x)$ denotes the logistic sigmoid transform which maps from the real number plane to a probability on the interval [0,1].

The authors analyzed the temporal cross-correlation of these scores and found significant evidence for the reinstatement of stimulus patterns in specific sequence ordering that matched the task structure; that is, scores followed in the patterns A->B->C->D and A'->B'->C'->D' (see Fig. 1B). This patterning was strongest for a time lag of $\tau = 40$msec, indicating very rapid serial reinstatement of visual stimulus representations. Finally, the authors estimated a single overall timecourse of replay $R_t$ using the following logical operation:

$$R_t = P\begin{pmatrix} (A_t \cap B_{t+\tau}) \cup (B_t \cap C_{t+\tau}) \cup (C_t \cap D_{t+\tau}) \\ \cup (A_t{}' \cap B_{t+\tau}{}') \cup (B_t{}' \cap C_{t+\tau}{}') \cup (C_t{}' \cap D_{t+\tau}{}') \end{pmatrix}$$

Where $\cap$ denotes the logical AND operation, and $\cup$ denotes the logical OR operation. Consequently, the replay timecourse, $R_t$, represents the probability of any task-relevant two item sequence occurring with the specified time lag $\tau = 40$msec. This probabilistic output was thresholded at the 99th percentile to provide the estimated replay event times used throughout this paper.

Resting State Network Modelling

The RSN-states referred to in Fig. 1 and throughout this paper use an established Hidden Markov Model with Time Delay Embedding (HMM-TDE) approach outlined in detail in *(15)*. We outline the approach here and direct readers to the original publication for further details.

This model makes use of the Hidden Markov Modelling framework, a generative modelling approach that describes the data $X_t$ at each timepoint $t$ as being generated from a latent state variable $Z_t \in [1,2, \ldots K]$. That is, the latent state at any point in time is an integer between 1 and $K$, where K is a parameter controlling the total number of states. Therefore, the DMN state being active at time $t$ would be denoted by $Z_t = 2$, with the DMN corresponding to RSN-state 2 in this case (Fig. 2B).

To complete the model specification, we must define the observation model – that is, the probabilistic model relating the activation of a particular latent state to the underlying data. As in *(15)*, we make use of a temporal embedding with a Gaussian observation model:

$$P(vec(X_{t-l:t+l}) | Z_t = k) \sim N(0, \Sigma_k)$$

In this notation, the $vec$ operator performs the temporal embedding; that is, we take a $[W \times P]$ matrix of points $X_{t-l:t+l}$ centered on the timepoint $t$, where P is the number of data dimensions and $W = 2l + 1$ is the length of the temporal embedding - and stack it into a $[WP \times 1]$ vector with the $vec$ operator. We then model this vector of datapoints with a Gaussian distribution, with zero mean and a $[WP \times WP]$ covariance matrix determined by the active state. Crucially, each RSN-state is now parameterized by a unique covariance matrix which reflects the autocovariance on each channel, as well as the cross covariance across channels. As outlined in *(15)*, this is an efficient parameterization of the power spectrum and the cross power spectrum respectively. Consequently, each distinct RSN-state corresponds to a distinct distribution of power on each channel and coherence between channels.

Model inference

The full model outlined above is amenable to variational Bayesian methods, which support fast scalable inference for large datasets. The datasets we have used are moderately large – the full model was trained on dataset C, comprising 5 minutes of resting state scans from 55 subjects at 250samples/second, totaling 3.9 million unique data samples; we then fit the learned model to datasets A and B, each comprising a total of 10 minutes of resting state scans from 21 and 22 subjects respectively, totaling 3.2 million unique samples and 3.7 million unique samples, respectively. To enable scaleable inference over such datasets, we utilized stochastic gradient variational Bayes methods that iteratively learn a full model through batch training, as outlined in *(46)*. Furthermore, we use PCA dimensionality reduction steps as outlined in *(15)* when training the model; to avoid potential misrepresentation of the data, the dimensionality reduced data is only used for model training, all spectral features plotted in figures are based on a multitaper fit back to the original, full dimensional data. Finally, hierarchical models fit with variational methods can be sensitive to local minima. To ensure good convergence, the full model inference was run five times; only the model with the lowest final free energy was kept, the others were discarded.

Model Parameter Selection

As a guiding principle we have maintained the same parameter choice as that previously used in *(15)*. The parameter $K$ controls the total number of RSN-states and therefore the granularity of the solution; we set this to $K = 12$, consistent with previous publications; the same results can be broadly reproduced with other choices of K.

RSN-state labelling

The HMM-TDE model infers a set of mutually exclusive latent states, which we number from 1 to 12 for ease of reference throughout. The RSN-state numbers themselves are arbitrary, however for ease of interpretation we assign labels using a data-driven approach that minimizes a distance metric between ordinal states. This can be visualized as in supplementary Fig. S1, and has the result that RSN-states that are nearer in ordinal label are considered nearer in the corresponding state space; for example RSN-state 2 can be considered nearer to RSN-states 1 or 3 than to RSN-state 12.

We derive this distance metric from the transition matrix directly. Let $\Theta$ be the Markov state transition matrix, such that each entry $\theta_{i,j} = P(z_{t+1} = j | z_t = i)$; that is, each entry contains the probability of observing a direct transition from state $i$ into state $j$. First, we exclude self-transitions, which are uninformative for the purposes of distances between states; this is done by setting the diagonal entries to zero, then normalizing so each row sums to one, such that our new matrix has entries $\psi_{ij} = P(z_{t+1} = j | z_t = i, z_{t+1} \neq i)$. Noting that high transition probabilities should correspond to small distances, we then define the distance between two states as the probability of not observing that state transition:

$$d_{ij} = 1 - \psi_{i,j}$$

Finally, to convert this to a symmetrical matrix of distances (as currently $d_{ij} \neq d_{ji}$), we simply average corresponding off-diagonal entries $\hat{d}_{ij} = \frac{1}{2}(d_{ij} + d_{ji})$.

Given the 12 x 12 matrix of distances $\hat{d}_{ij}$, we then use multidimensional scaling to identify the single axis that accounts for the most variance in the distance matrix. RSN-states are then labelled 1-12 in the order of their appearance on this axis – see visualization in Fig. S1.

RSN Spectral Information

Each RSN-state is defined by a distinct spatial distribution of spectral power and coherence. These characteristics can be interpreted directly from the observation model parameters - however this may emphasize certain traits of the model fitting procedure, in particular the effects of the PCA dimensionality reduction methods applied, rather than the underlying spectra that is observable in each RSN-state. For this reason, we extract the spectral information by fitting a multitaper to the raw data, conditioned on the active RSN-state as introduced by *(47)* which provides an empirical assessment of the power and coherence for each ROI as a function of frequency. As outlined above, due to the increased confidence of spatial RSN-state parameters derived from dataset C, a larger dataset for which MRI structural information was available, the spatial and spectral distributions per RSN-state referred to in this paper were derived from this dataset, supporting their interpretation as a set of canonical RSN-states.

This information is still very high dimensional so we summarize it through a spectral mode decomposition, obtaining spatial power and coherence maps for a data-driven set of frequency band modes. This decomposition is implemented by non-negative matrix factorization as detailed in *(15)*. For summary purposes in Fig. 2B, Fig. S2B and Fig. S3, we fit this with two modes, to separate a single wideband mode from higher frequency noise. To explore further the spatial breakdown of different frequency modes, we fit the same non-negative matrix factorization procedure with four modes as in Fig. 5 (use of four modes was found to produce broadly stable decompositions and mirrors findings in previous work). These can be derived for each RSN-state; alternatively, to more accurately reflect the full power and coherence distribution at a particular event time, we can use the evoked RSN-state distribution to weight the power and coherence information, producing HMM regularized time frequency plots in Fig. 5 and Fig. S5.

Replay Evoked RSN Analysis

Fig. 2A analyses the average RSN-state distribution evoked by individual replay events. At the single subject level, we took the probabilistic RSN-state timecourse $\{\gamma_{i,t}\} = P(Z_t = i)$; that is, a timeseries with K distinct values between 0 and 1 at each timestep reflecting the probability of each RSN-state's activation. We then baseline corrected by subtracting the average over all timepoints for that subject, such that values greater than zero reflect probabilities greater than the session average, and values less than zero reflect activation probabilities below the session average for that subject. We then epoched this time series using the replay times identified above, extracting a window extending from 0.5 seconds prior to each event to 0.5 seconds after each event. For each subject, we then computed the evoked RSN-state distribution over this time window by averaging the epoched RSN-state distributions over all replay events. Let us denote by $B_{i,\tau,n}$ the evoked distribution for subject $n$ and RSN-state $i$ at timelag $\tau$ from the estimated replay events. Fig. 2A plots the mean of these values over subjects, reflecting the expected increase or decrease in RSN-state probability evoked by individual replay events.

To test the significance of these results two types of statistical test were utilized; to support claims of a significant result at a specific point in time we used a one-sided t-test, whereas to support claims of a broad peak of significant points across time we used a non-parametric sign flipping cluster permutation test. The cluster permutation test determined the probability of finding by chance a cluster of consecutive significant points that was the same size or larger than that observed in the data. We defined significance in this context as observing a group level t-statistic exceeding 3; we then generated a null distribution of cluster sizes by randomly permuting 5000 times the sign of each subject's evoked response parameters $B_{i,\tau,n}$ before computing group t-statistics, and determining the number of consecutive points exceeding the significance threshold. We could thus compute the chance probability of generating a cluster of the same size observed in the data.

Fig. 2C plots the results of the same analysis on the functional localizer data; epochs were defined by the time of visual stimulus presentation, corresponding to t=0 in the plot. The thick red line indicates the time at which the replay classifiers were trained.

Fig. 2D then directly compares the evoked RSN-state distribution between the replay and functional localizer data sessions; for ease of presentation, we only focused on the distribution at the exact time of replay onset (t=0) and the exact timepoint the replay classifiers were trained on (t=200msec); for each of the 12 RSN-states we conducted a paired t-test to check whether the evoked RSN-state distributions were significantly different across the two conditions.

Transient Burst Activity Analysis

To analyze the temporal statistics of replay onset, we modelled the estimated replay event times for each subject and as a Poisson point process. Wherever an interval between events encompassed MEG data samples that had been marked as bad, the whole interval was excluded. We then partitioned the replay timecourse into non-overlapping windows, for a given window length W, and computed the total sum of replay events observed in each window *(48)*. The Fano Factor, for a given window of length W, is then given by:

$$F_W = \frac{\sigma_W^2}{\mu_W}$$

Where $\mu_W$ and $\sigma_W^2$ are the mean and variance of event counts in each window. The Fano Factor was computed individually for each subject, Fig. 3D plots the mean +/- ste over subjects for each window length. We then computed parametric t tests on the Fano factor to test the null hypothesis that it was equal to one.

This alone does not prove conclusively that replay occurrences are transient burst events, but rather that they are not homogenous Poisson processes. Hence, we furthermore tested a much broader null hypothesis that the intervals between replay events were independent and identically distributed, following the approach of *(48)*. We generated surrogate data by shuffling the interval times; that is, for each subject, we took the replay event timecourse, computed the intervals between consecutive events, and randomly permuted the intervals to generate a new event timecourse. This maintained the exact distribution of replay intervals whilst removing any dependence between consecutive intervals. We then computed the subject and group Fano Factors exactly as above. With this surrogate data, after 1000 permutations the maximum Fano Factor computed (plotted as the dotted line in Fig. 3D) remained well below that of the true timecourse, confirming that the longer term dispersion arises from correlation over consecutive intervals ($p<1e-3$) and allowing us to conclude that replay is characterized by irregular transient bursting behavior.

We then repeated the analysis for RSN-state activations (Fig. 3E). Treating each RSN-state visit as an event, we removed consecutive activations and just used the interval between events to create the equivalent RSN-state visit event timecourse. As above, we computed the Fano Factor vs window length for each subject.

To compute the significance of these results, we used a one-way ANOVA to test the null hypothesis that there was no significant difference between the Fano factors of different RSN-states; this was conducted for each window length resulting in 100 multiple ANOVA tests. We only report the highest p value obtained, which remained significant with Bonferroni correction, allowing us to reject the null hypothesis for all window lengths observed. We could then assess a two-sample t-test, taking each RSN-state one by one and testing the hypothesis that the observed Fano Factors for that RSN-state were no different from the entire population of remaining RSN-states. For example, we would take the 21 subject level Fano Factors for RSN-state 1, and compare that with the population of size 21 by 11 of Fano Factors for each subject for RSN-states 2-12, then repeat this for RSN-state 2. Again, this test was computed for each value of the window size; only the highest obtained p value was reported, all others were below this and therefore more significant.

Finally, Fig. 3F considers the interval between replay events given the active RSN-state; that is, if a replay event occurs at time t we compute the length of time to the successive replay event, and also the most likely RSN-state to be active at time t. In this analysis, we omit any observations where the interval to the next replay event overlaps with segments marked as bad. For each subject, we thus compute a mean replay interval given the active RSN-state, generating a collection of 21 observations of replay intervals given the active RSN-state. Certain subjects had very few observed replay events in particular RSN-states, resulting in high variance estimates; we therefore omitted from further analysis any means that were derived from fewer than ten observations. We then could then test for significant variation by active RSN-state with a one-way ANOVA followed by two sample t-tests, in which we tested whether the replay intervals given RSN-state k was active were significantly different from the replay intervals of all other RSN-states combined.

RSN-state Power and Coherence Analysis

Fig. S3 is another way to characterize the spectral information unique to each RSN-state. For these plots, we take the RSN-state spectral information as outlined by the multitaper approach outlined above.

To summarize the wideband response over all frequencies, we take the NNMF decomposition with two modes – that is, one mode that we interpret as wideband and another we interpret as higher frequency noise – and plot the RSN-state specific power and coherence for each ROI, given the active RSN-state. The power is given directly by the multitaper approach; coherence is defined per pair of ROIs, so the values plotted on the y axis in these plots is the sum of coherences between a given ROI and all others.

We can also follow the same procedure for each of the three spectral decompositions corresponding to individual spectral modes; that is, we derive the power and coherence for each ROI and for each given RSN-state, as computed for each of the delta / theta, alpha and beta spectral modes.

Replay Evoked Spatial Maps

To accurately convey the spectral profile that is linked to each replay event, we can combine the replay evoked RSN-state distribution plotted in Fig.2A with each RSN-state's spectral information. As outlined above, we have values for the spectral power and coherence of each RSN-state from dataset C. We can initially visualize these by averaging over all recorded ROIs; the left-hand side of Fig. 5A and Fig. 5B plot the state specific power and coherence values, respectively. To compute the power and coherence patterns that most closely reflect replay times, we take the subject specific replay evoked RSN-state distributions, referred to as $B_{i,\tau,n}$ above and plotted in the lower panel of Fig. 5A and Fig. 5B. For consistency of interpretation with Fig. 2 these are plotted after baseline correction, however to compute the replay evoked power and spectral density plots we revert to the non-baseline corrected values, such that each $B_{i,\tau,n}$ is a probability between 0 and 1, with the sum of values over all different states equal to 1 for each timepoint and subject. We can then compute the expected power and coherence as a function of both time and frequency around each replay event by weighting the state specific PSD and coherence values by the replay evoked state distribution. Note that this analysis was done at the subject level; that is, we computed subject-specific power and coherence estimates as a function of both time and frequency using each subject's specific replay evoked state distribution. This produced maps of dimension [time x frequency x subjects x channels] in the case of PSD and [time x frequency x subjects x channels x channels] in the case of coherence. Given the high dimensionality of this data we separately summarize the time-frequency information (Fig. 5A and Fig. 5B) and the spatial frequency distribution (figure 5c). For visualization in the heatmaps of Fig. 5A and Fig. 5B we baseline corrected at the subject level, such that zero on the color scale denotes an average PSD and coherence. Fig. 5C then expresses how this information is distributed over frequency and space. This plot uses only the distribution defined at t=0, the exact time of replay estimated by Liu et. al. (16). We decompose the spectral information using the previously introduced non-negative matrix factorization, a data-driven approach that identifies the main spectral modes explaining the data, producing a vector of power values over channels and a matrix of coherence values between channels for each spectral mode and each subject. For the purposes of visualization, PSD values are thresholded to plot the highest 50% of values. The higher dimensionality of the coherence matrix required a more stringent thresholding approach; for this we applied a gaussian mixture model threshold test *(15)*, only plotting connections if they fall into a distinct mixture separated from the global distribution. All these spectral estimation and thresholding steps follow those previously introduced *(15)*.

<u>High Frequency Oscillations</u>

To analyze the higher frequency correlates of specific RSN-state activations, we first took the timeseries of state activations that had been fit to the resting state data as outlined above. Importantly, this analysis had been conducted on data that was downsampled to 250 samples/second and band pass filtered between 1 and 45Hz. For this analysis we therefore returned to the raw data which had been acquired at 600samples/second with a low pass filter applied with cutoff 160Hz. This data did contain power line noise with a baseline frequency at 50Hz and some harmonic noise at integer multiples of 50Hz; we did not attempt to filter this out to avoid introducing any undesirable filter effects. Using the exact state activation times and multitaper approach outlined above, but instead using data with a higher sampling rate that included high frequency content, we estimated the power spectral density associated with each state over all frequencies from 1 to 160Hz. Fig. 6A plots the mean +/- standard error over subjects of the PSD across this broader frequency band for all states. To compare this to the PSD estimated at replay times, we reused the same multitaper approach but replaced the state activation times with estimated replay times. Specifically, we defined a window of width 20msec around the time of each estimated replay event, and computed the PSD across all frequencies for the data in these windows using the same multitaper approach; we furthermore computed the PSD for all timepoints excluding these replay times to use as the baseline. These PSD values are plotted in Fig. 6B. Finally, given the exclusive correlation between high frequency power increases and RSN-state 2, we explored the spatial distribution of power in higher frequencies in this specific RSN-state. Fig. 6C averages the PSD values over all subjects between 102-148Hz (this band was defined to exclude mainline power noise effects) for each ROI, and plots the spatial topography over all ROIs.

#### Supplementary Text

Resting State Networks in MEG and fMRI

MEG and fMRI are highly complementary modalities for imaging brain function, with MEG offering enhanced temporal resolution with courser spatial resolution, whilst fMRI offers high spatial resolution and poorer temporal resolution. Computational methods allow resting state networks to be derived in either modality to take advantage of the spatial / temporal resolution tradeoffs, however given the fundamentally different physiological basis of the MEG and fMRI signals, we are not afforded a straightforward correspondence between the inferred networks. We here summarize existing results as justification of our claims throughout the paper regarding how the MEG-derived RSN-states relate to canonical RSNs more typically identified in fMRI, particularly with regard to the DMN and DAN networks.

We have identified RSN-state 2 as the DMN state on the basis of its spatial, temporal and spectral properties, which are consistent with existing MEG literature and have been shown in simultaneous EEG/fMRI recordings to correlate with the fMRI DMN. The DMN was originally identified in fMRI and is taken to include the medial prefrontal cortex, posterior cingulate cortex and lateral parietal cortex as core regions; additional areas including the orbital frontal cortex, entorhinal/parahippocampal cortex and lateral temporal cortex have moderate correlations with this network leading to their intermittent inclusion in the DMN *(23)*. In MEG, activity patterns consistently replicate this anatomical distribution subject to the reduced spatial resolution the method provides (in particular, higher variance has been noted over cingulate regions - MEG has reduced sensitivity to deep cortical surfaces) *(10,13)*. RSN-state 2 extracts broadband patterns of activity in parietal cingulate and lateral parietal areas; upon analyzing the narrowband signal this also identifies frontal, anterior cingulate and lateral temporal areas active in a lower frequency band to the dominant alpha band. This is consistent with the previous MEG literature on the DMN *(15)*. Most importantly, this network matches very closely that identified in simultaneous EEG and fMRI recordings that correlates with the fMRI DMN network *(20,21)*. Furthermore, beyond the spatial correlation, this network similarly replicates key spectral and temporal properties of the DMN inferred in previous MEG literature; namely an increase in power *(14)* and coherence *(15)*, a relatively low fractional occupancy, or time active in the recording *(14,15)* combined with longer interval times and longer state lifetimes than other states *(14,15)*.

We have identified RSN-state 1 as the Parietal Alpha network, and posit that this is the equivalent of the DAN switching off. This is on the basis of this RSN-states spectral decomposition, which identifies a prominent alpha mode over parietal areas (Fig. S7). The spatial topography of the DAN in fMRI consists of activity in inferior parietal sulcus, superior parietal lobule and frontal eye fields *(49)*; the poorer spatial resolution of MEG cannot discern these regions individually, but identifies periods of increased activity over broadly parietal regions in the alpha band as the MEG DAN equivalent *(13)*. Most importantly, simultaneous

EEG/fMRI experiments have found that that the EEG equivalent to the MEG-derived RSN-state 1, anticorrelates with the fMRI DAN *(20,21)*. This supports our interpretation that activation of RSN-state 2, the parietal alpha network, signifies the DAN switching off, which is similarly believed to represent the inward direction of attention.

Controlling for Replay Classifier Variance During DMN Activation

RSN-states 1 and 2, which correlate with replay, also have the highest variance in the data (as in Fig 4, these RSN states have the highest broadband power and therefore the highest variance). This leads to increased variance in the classifier outputs when these states are on (Fig. S8A), which could lead to spurious "reactivations" occurring solely as a result of the increased variance in the classifier outputs.

To explore this further, we assert that such spurious effects must be unbiased; that is they should not selectively bias a classifier output towards a particular value. It therefore follows that, when classifier outputs are plotted by percentile, any spurious effects attributed to increased variance should be symmetric across the median. In Fig. S8B we plot the RSN-state distribution evoked by different percentiles of the replay score. This plot shows that the association with RSN-states 1 and 2 reported in Fig. 2A is robust over a broad range of percentile thresholds. In the corresponding lowest percentile range, a small effect does indeed emerge for the lowest 1 percentile, suggesting that the classifier variance does in some way contribute to the RSN-state replay relationship. This effect however is not significant after Bonferroni correction, and is only a fraction of the effect size in the highest percentile (Fig. S8C and S8D). Finally, we can illustrate that the result of Fig. 2A is robust over controls for this effect by plotting instead the contrast between $99^{th}$ and the $1^{st}$ percentile, as in Fig. S8E; the key result remains significant ($p<2e-4$ cluster significance test).

Replay Evoked and Reactivation Evoked State Distributions

Neuroscience recognizes two distinct but closely related phenomena occurring in the brain. Reactivation refers to the reinstatement of neural traces of recently acquired information; replay is the serial reactivation of multiple such traces in a temporally compressed manner *(6-7)*. Consequently, reactivation is a requirement for replay to be detected, but not all reactivation constitutes replay. An important question that follows from our work is whether the relationships we have observed are true for reactivations in general, or whether they are preferentially true for reactivations occurring in replay sequences. We find no evidence of distinct relationships between RSN-states and reactivation as opposed to replay, however also caution against any strong interpretation of this result given the limitations of our methods outlined below.

Fig. S9A replicates the replay evoked RSN-state distribution of Fig 2. Fig S9B conducts the same evoked response analysis, but instead of using the timecourse of replay, utilizes the timecourse of reactivations – that is, the actual stimulus classifier outputs denoted by $, A_t, B_t, \ldots$ in the methods section on Replay Detection. The result is a very similar evoked distribution.

However, the methods we have used here are not well suited to discern differences in evoked distributions between RSN-states and replay vs reactivation. Following Liu et. al., we have defined replay as the linear superposition of reactivations with a given timelag; given the autocorrelation structure of the reactivation timecourses this linear superposition remains closely correlated with the overall reactivation timecourse (Pearson's correlation coefficient r=0.45). Furthermore, given the bursty nature of replay that we have demonstrated and the corresponding temporal irregularity (bursty nature) of the RSN-states, the similarity of the evoked RSN-state distributions shown in Fig S9B is perhaps unsurprising. It remains possible that underlying differences between RSN-state activity associated with replay and reactivation do indeed exist, however, the existing methods are not sensitive to them.

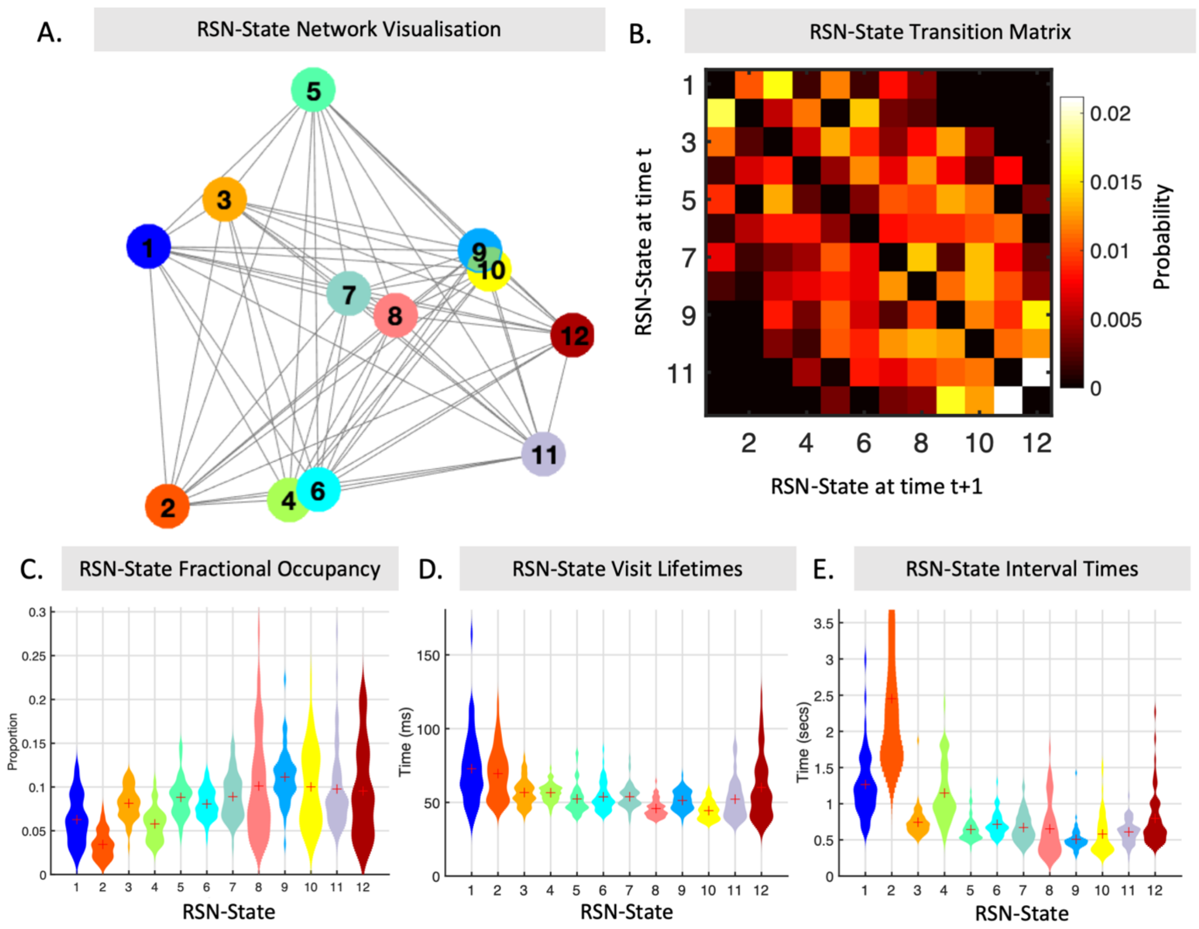


**Fig. S1.**

Resting State Network Temporal Profiles. (A) The RSN-states can be visualized as a full network through a multidimensional scaling of the transition matrix (see Methods). We label RSN-states 1-12 in order of their appearance on the horizontal axis of this graph, accounting for the greatest portion of distances between RSN-states. (B) The transition matrix (excluding diagonal entries) for the RSN-states; the diagonal structure reflects this labelling structure, with transitions more probable between adjacent pairs of RSN-states than distant pairs. (C) RSN-state fractional occupancies; the proportion of total timepoints that are occupied by each state; violin plot shows distribution over subjects, with crosses denoting mean. RSN-states 1 and 2 explain proportionally less of the data than the remaining RSN-states. (D) RSN state visit lifetimes; the average length of a visit to each RSN-state; violoin plots show distribution over subjects, crosses denote mean. When visited, RSN-states 1 and 2 are activated for longer than any other RSN-states. (E) RSN-state Interval times; the average length of time between consecutive RSN-state visits. RSN-states 1 and 2 display longer interval times than other RSN-states.

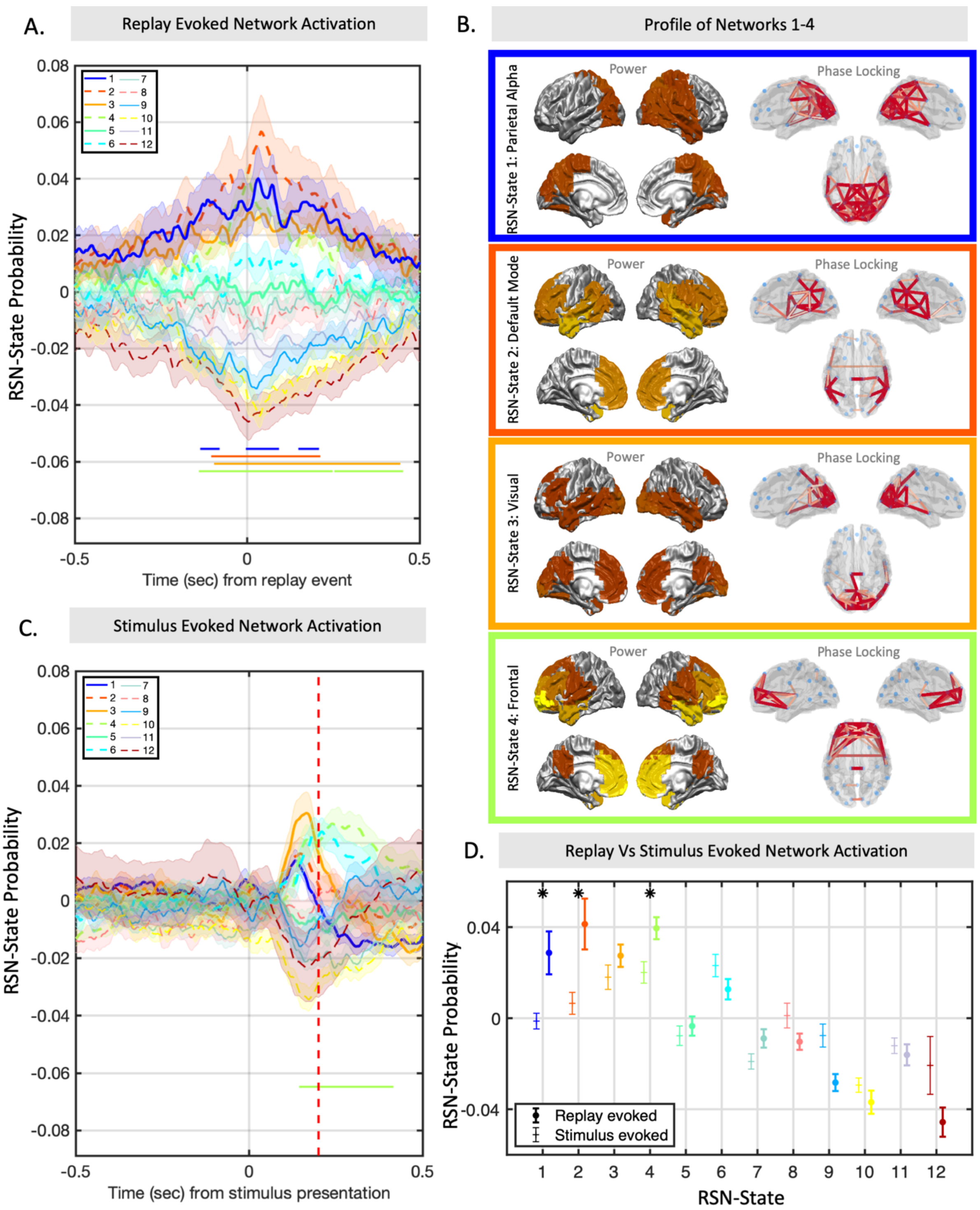


**Fig. S2.**

Replicating the result of Fig. 2 on a second dataset. (A) Plotting mean +/- ste of the RSN-state probability evoked by replay onset; replay onset coincides with a broad peak in activity in RSN-states 1 and 2, and to a lesser extent RSN-states 3 and 4. Significance bars show clusters where p<2e-4. (B) Wideband activity profiles for RSN-states 1-4 (see Methods). (C) Mean +/- ste of the

RSN-state probability associated with the original stimulus data. Significance bars show clusters where $p<2e-4$. (D) Comparing directly the mean +/- ste of the evoked state distribution at replay time and at the classifier training time identifies RSN-states 1 and 2 as significantly increased during spontaneous replay (multiple paired t tests). Single asterisk denotes $p<0.05$, double asterisk denotes $p<5e-4$. Inset: Result of replication study on second dataset.

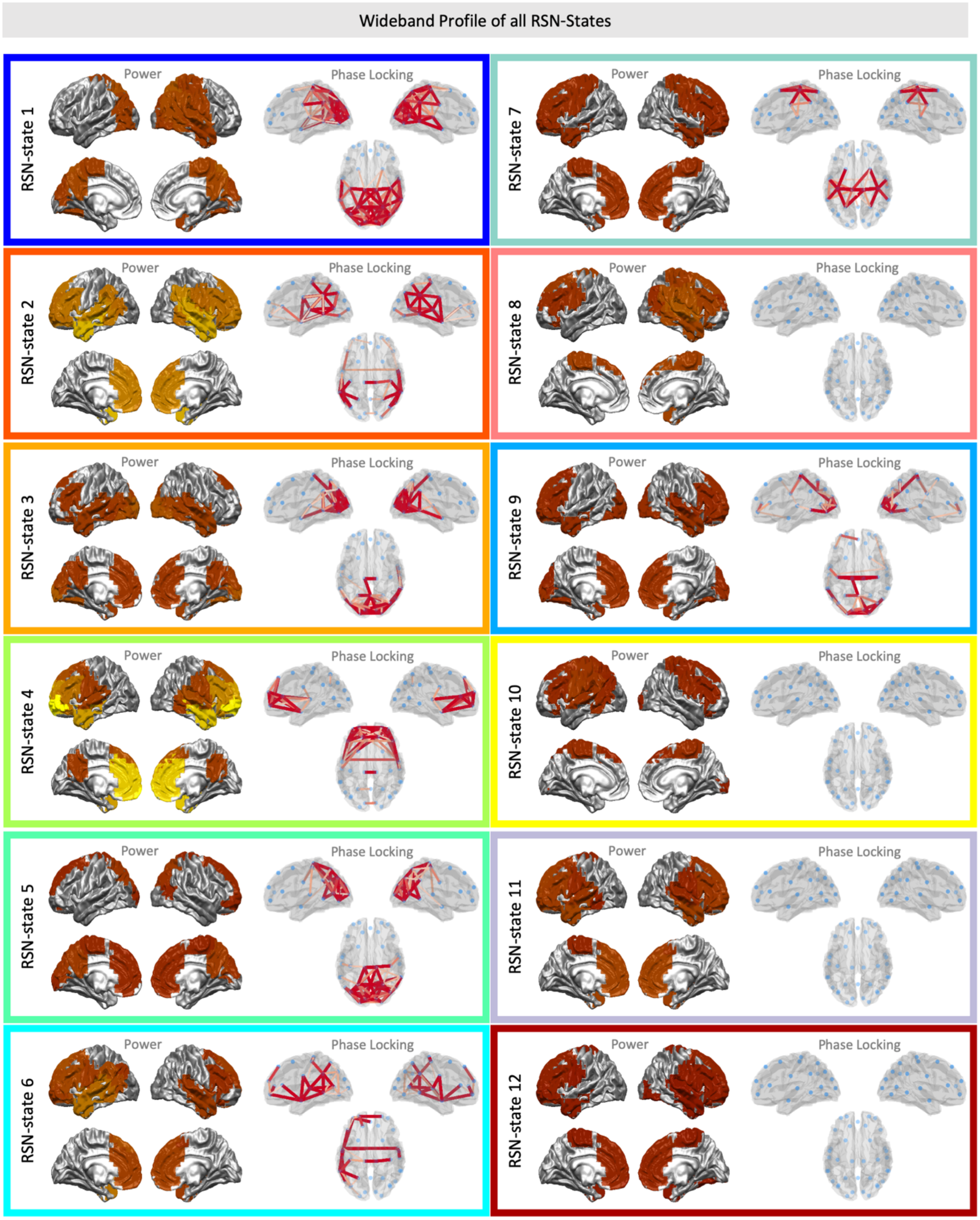


**Fig. S3.**

Wideband profiles for all RSN-states, plotting the wideband power (thresholded at 50%) and coherence (thresholded with Gaussian Mixture Model). See Methods for further details and Supplementary Text for clarification of how these relate to equivalent networks in fMRI.

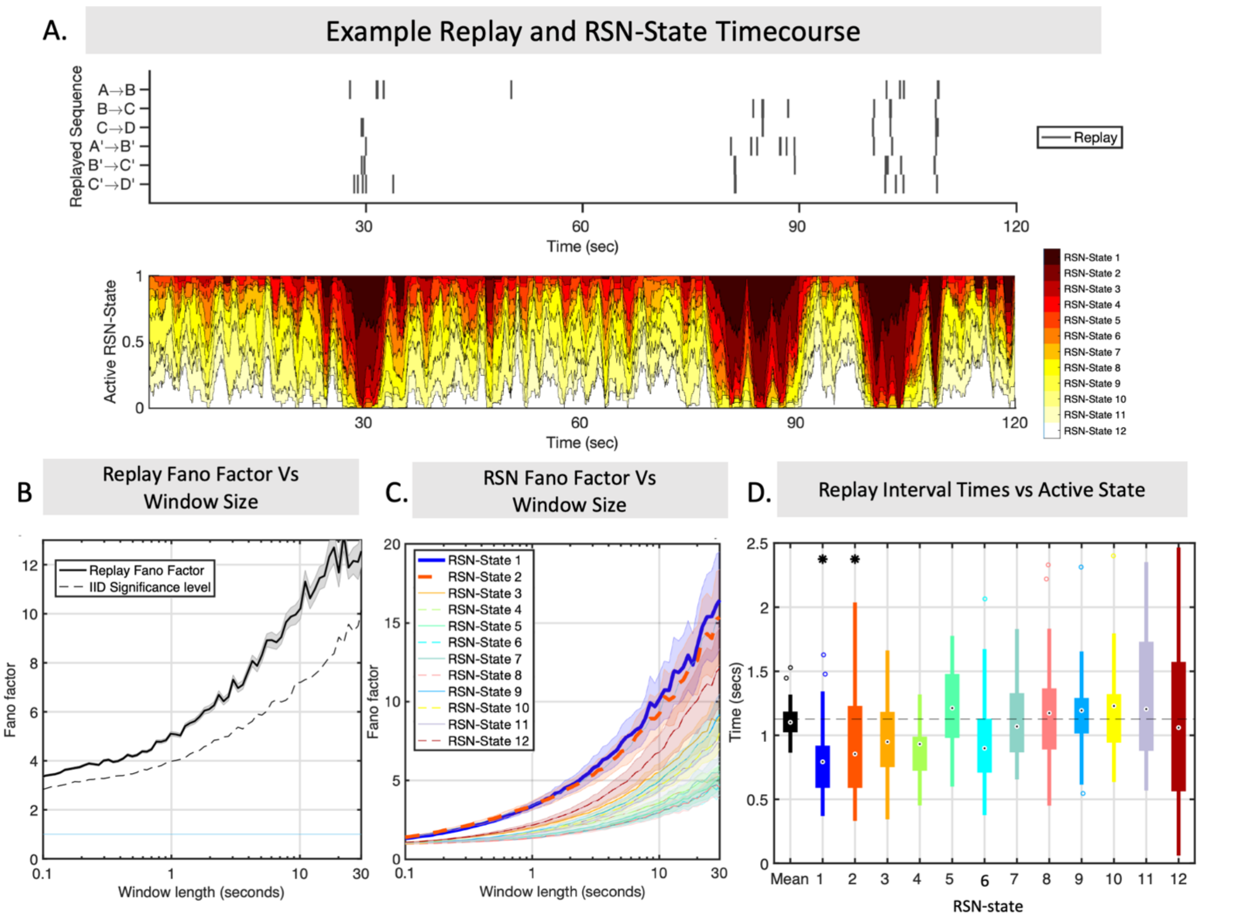


**Fig. S4.**

Replication of Fig. 3 results on a second dataset. (A) Example subject data showing the bursty nature of replay coinciding with relatively infrequent visits to the replay-associated RSN-states. For ease of visualization, lower panel plots the one second moving average RSN-state probability. (B) Temporal irregularity can be quantified by the Fano factor as a function of window size. Replay events show that this irregularity measure increases over longer timescales, displaying maximum temporal irregularity over windows of ten seconds or more. (C) This structure is replicated by the RSN-state activations, with RSN-states 1 and 2 displaying the most irregular patterns at long timescales. (D) This temporal structure is not just common but in fact coincides; replay events that occur during RSN-state 1 or 2 have significantly shorter periods, reflecting rapid bursty behavior during the infrequent state visits and long periods of quiescence outside of these.

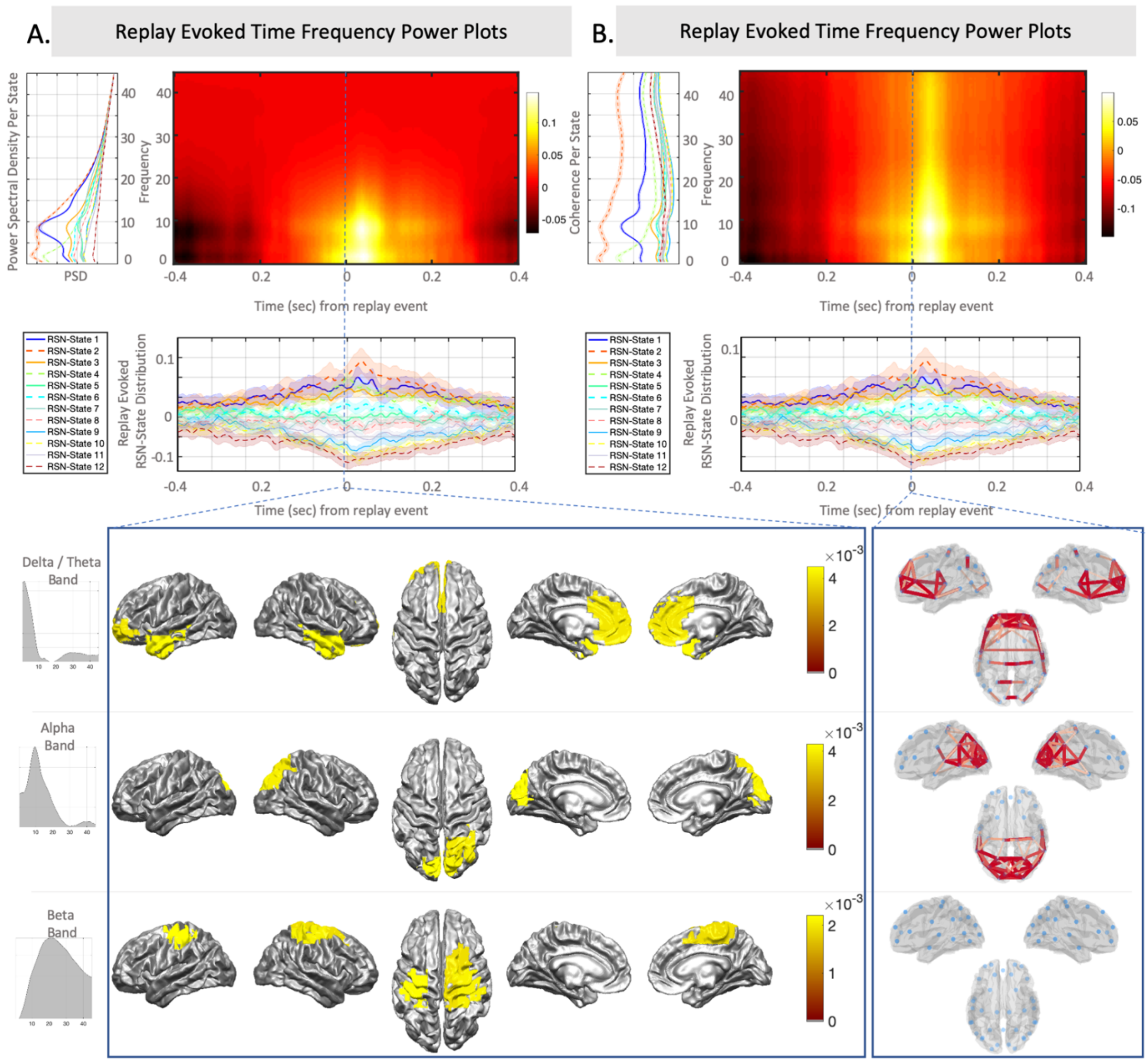


**Fig. S5.**

Replication of results in Fig. 5 on a second dataset. (A) We can use the PSD estimates of each RSN-state (left) and the replay evoked RSN-state probabilities (lower) to reconstruct a time-frequency estimate of power spectral density around replay events, revealing a prominent peak in the alpha and delta /theta bands. (B) In addition to increases in power, the replay associated RSN-states show an increase in coherence across all frequencies, but especially in the alpha and delta/theta bands. (c) Plotting the spatial distribution of activity in the defined frequency modes at the time of replay identifies independent modes of coherent activity; a low frequency mode comprising frontal DMN and temporal areas, and an alpha frequency mode comprising parietal DMN regions and visual cortex. Additionally, some weaker levels of activity in the beta band are observed over motor areas, but network coherence patterns in this frequency band are not significant.

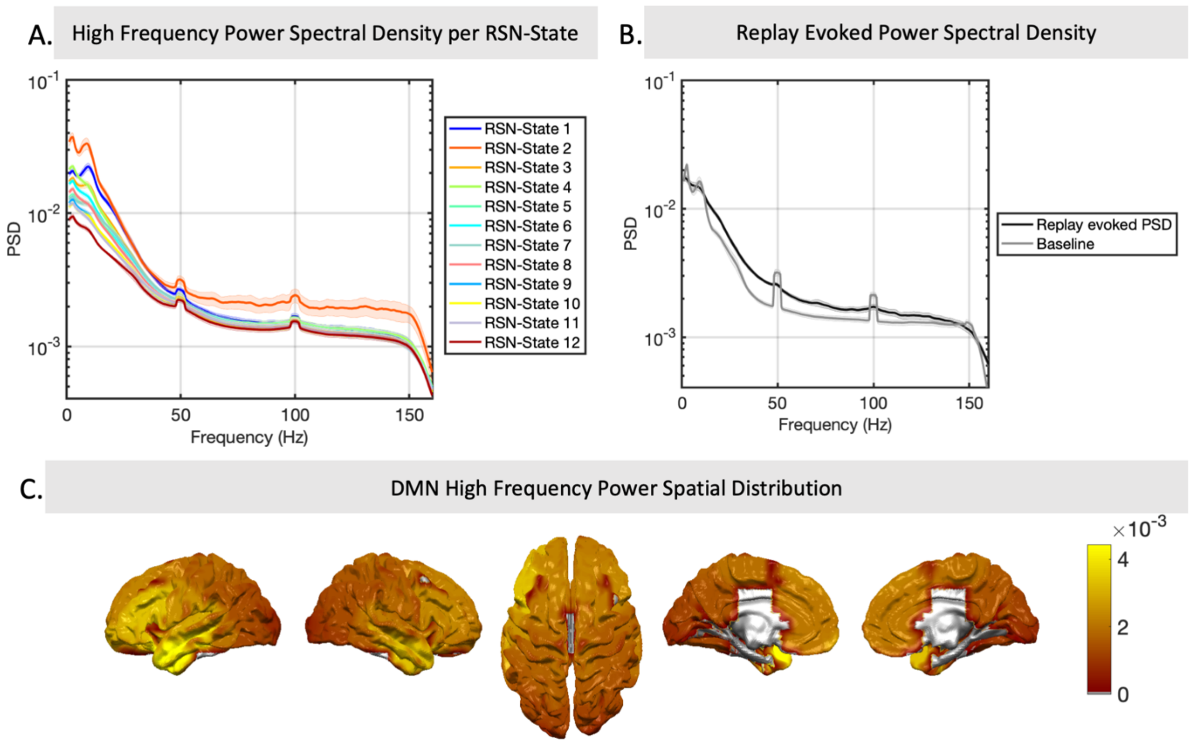


**Fig. S6.**

Replication of the result of Fig 6 on a second dataset. (A) Although the RSN-state model was originally fit to data filtered at 1-45Hz, we can still analyze whether the state timings correlate with specific patterns in frequency bands outside this range in the original data. This reveals a very strong association between RSN-state 2 and power in high frequencies – despite these high frequencies not having been originally included in the model. (B) Similarly, the onset of replay is associated with an increase in high frequency power relative to the global average. (C) Activity in this RSN-state and in this frequency band may originate in temporal cortices.

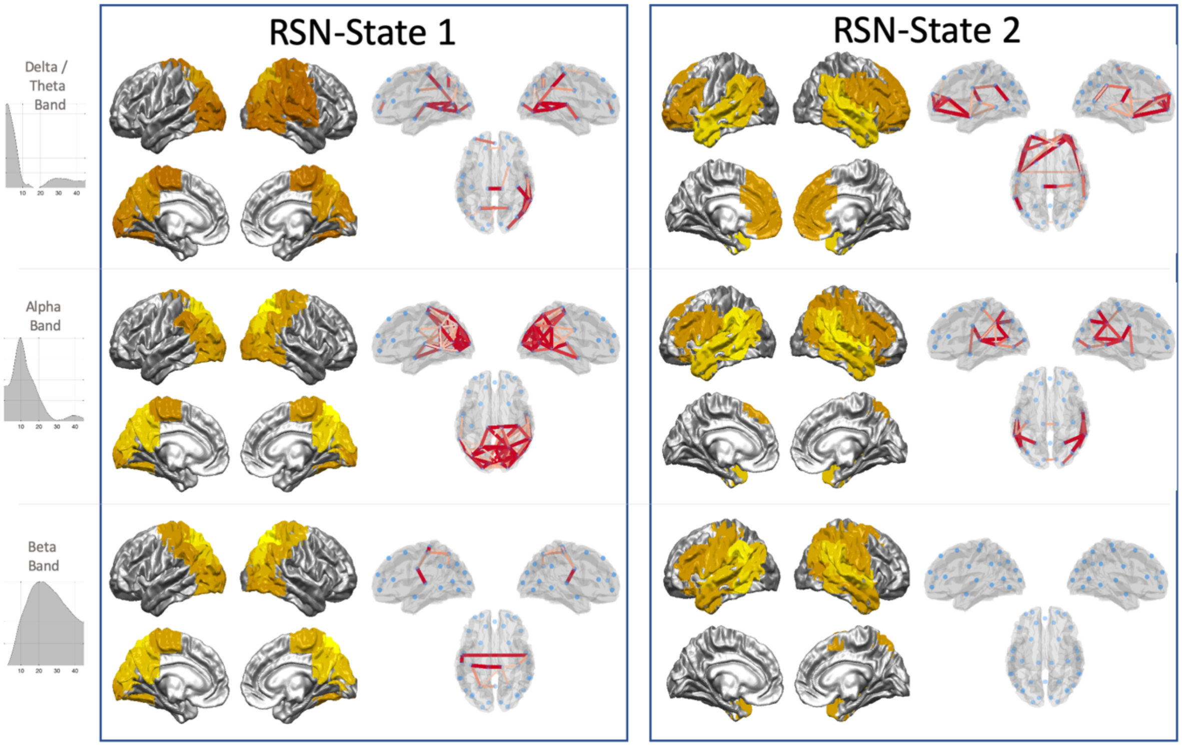


**Fig. S7.**

The full spectro-spatial activity maps associated with RSN-states 1 and 2; where Fig. S3 plotted the wideband distribution of power, this can be further analyzed using a data driven spectral decomposition. We here plot the spatial distribution of power in each frequency band (thresholded at 50%); along with the coherence network (Gaussian mixture model threshold; see Methods). This supports our interpretation of RSN-state 1 as a parietal alpha state, given the dominance of alpha band power and coherence over parietal cortex. Similarly RSN-state 2 breaks into the frontotemporal delta/theta band mode and lateral parietal alpha band coherence that associated with the DMN in MEG.

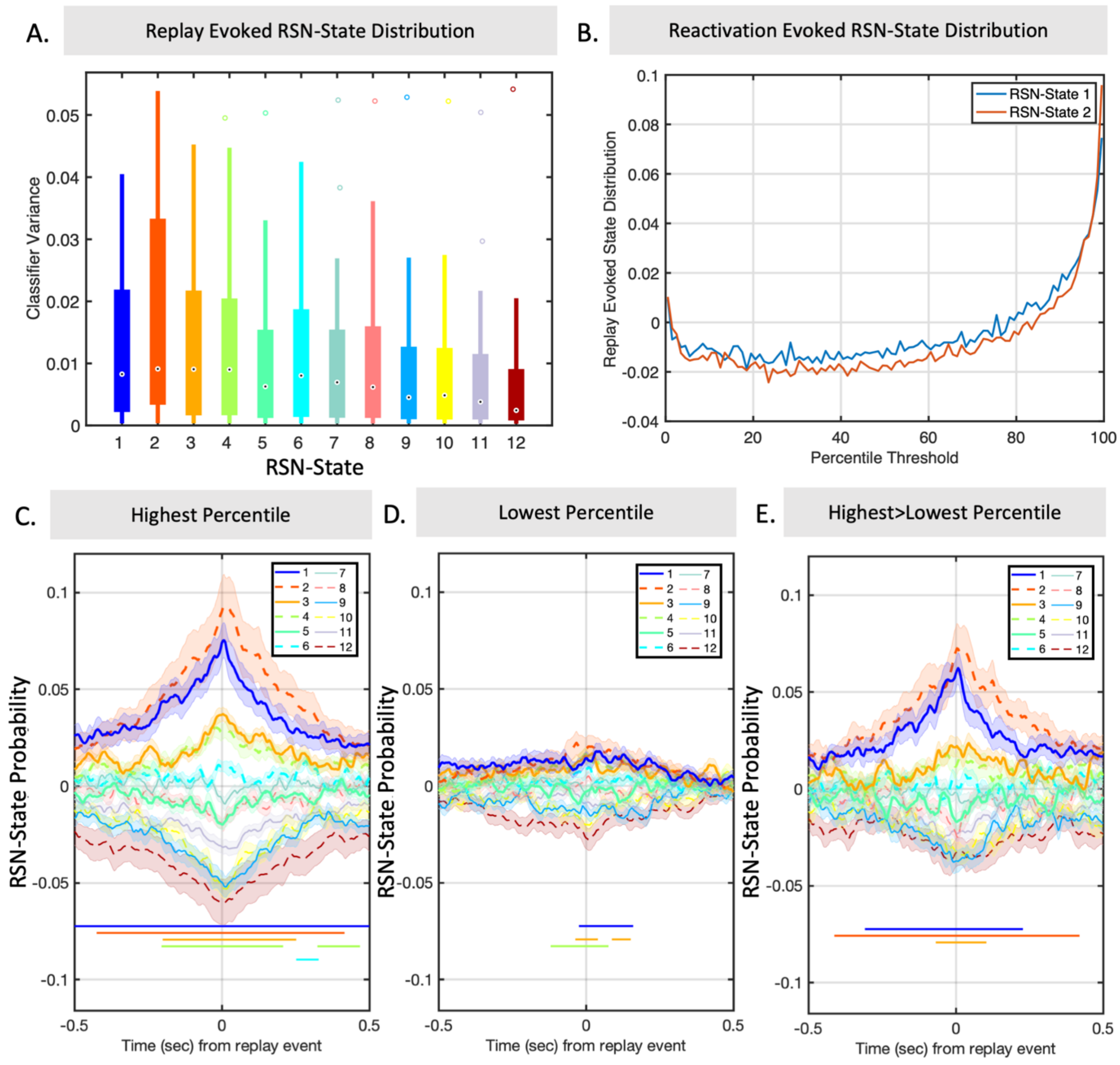


**Fig. S8.**

Controlling for RSN-state specific increases in classifier variance. (A) Variance of classifier output varies as a function of RSN-state (One way ANOVA, $p < 0.01$). Box plots show variation over subjects of observed classifier variance during activation of each RSN-state. (B) Focusing only on RSN-states 1 and 2, the replay evoked state distribution at $t=0$ (ie the exact time of estimated replay onset) is plotted against the percentile threshold used. This identifies a relationship that is not monotonically increasing, but rather has two peaks at the highest and lowest threshold, reflecting that some of the RSN-state correlation derives from the increased variance of the classifiers when RSN-states 1 and 2 are active. (C) Replicating the main result of Fig. 2A, the RSN-state distribution evoked by the highest percentile of replay scores, for comparison. (D) The RSN-state distribution evoked by the lowest percentile of replay scores. (E) Taking the contrast of these as a control for the effect size that is solely due to classifier variance again replicates a strong association between RSN-states 1 and 2 and the onset of replay.

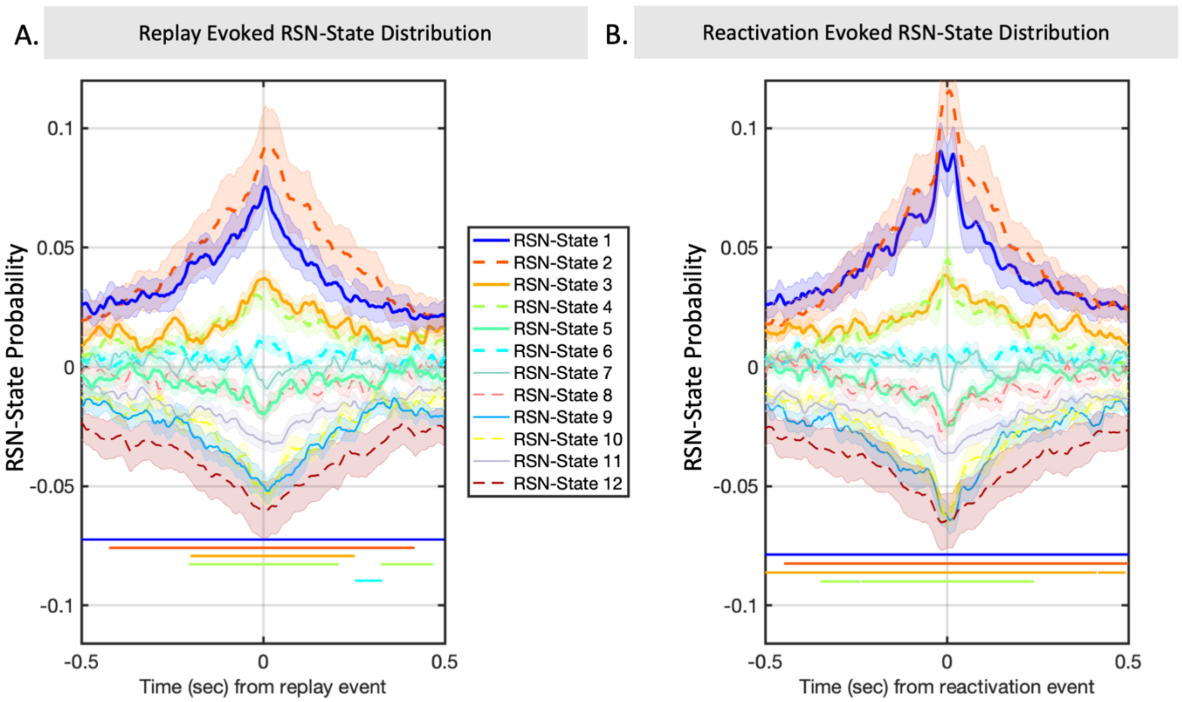


**Fig. S9.**
The replay evoked RSN-state distribution is a special case of the reactivation evoked RSN-state distribution. (A) The main result of Fig. 2A, reproduced for ease of comparison. (B) The corresponding RSN-state distribution evoked by reactivation (see Methods and Supplementary Text). Significance bars denote clusters with $p<2e-4$.
